# A putative *de novo* evolved gene required for spermatid chromatin condensation in *Drosophila melanogaster*

**DOI:** 10.1101/2021.06.10.447990

**Authors:** Emily L. Rivard, Andrew G. Ludwig, Prajal H. Patel, Anna Grandchamp, Sarah E. Arnold, Alina Berger, Emilie M. Scott, Brendan J. Kelly, Grace C. Mascha, Erich Bornberg-Bauer, Geoffrey D. Findlay

**Affiliations:** Department of Biology, College of the Holy Cross, Worcester, MA, USA; Institute for Evolution and Biodiversity, University of Münster, Münster, Germany; Department of Protein Evolution, Max Planck Institute for Developmental Biology, Tübingen, Germany; Department of Molecular and Cellular Biology, Harvard University, Cambridge, MA, USA; Department of Organismic and Evolutionary Biology, Harvard University, Cambridge, MA, USA; Department of Entomology, The Ohio State University, Columbus, OH, USA

## Abstract

Comparative genomics has enabled the identification of genes that potentially evolved *de novo* from non-coding sequences. Many such genes are expressed in male reproductive tissues, but their functions remain poorly understood. To address this, we conducted a functional genetic screen of over 40 putative *de novo* genes with testis-enriched expression in *Drosophila melanogaster* and identified one gene, *atlas,* required for male fertility. Detailed genetic and cytological analyses showed that *atlas* is required for proper chromatin condensation during the final stages of spermatogenesis. Atlas protein is expressed in spermatid nuclei and facilitates the transition from histone- to protamine-based chromatin packaging. Complementary evolutionary analyses revealed the complex evolutionary history of *atlas*. The protein-coding portion of the gene likely arose at the base of the *Drosophila* genus on the X chromosome but was unlikely to be essential, as it was then lost in several independent lineages. Within the last ∼15 million years, however, the gene moved to an autosome, where it fused with a conserved non-coding RNA and evolved a non-redundant role in male fertility. Altogether, this study provides insight into the integration of novel genes into biological processes, the links between genomic innovation and functional evolution, and the genetic control of a fundamental developmental process, gametogenesis.

**Author Summary:** Genomes are in flux, as genes are constantly added and lost throughout evolution. New genes were once thought to arise almost exclusively via the modification or duplication of existing genes. Recently, however, interest has grown in alternative modes of new gene origination, such as *de novo* evolution from genetic material that previously did not encode proteins. Many *de novo* genes are expressed in male reproductive tissues, but their significance for fertility is not well understood. We screened likely *de novo* genes expressed in the *Drosophila* testis for reproductive roles and found one gene, *atlas*, essential for male fertility. We leveraged genetic and cell biological experiments to investigate roles for Atlas protein in reproduction and found that it is required during sperm development for proper packaging of DNA in the sperm nucleus. Evolutionary analyses of this gene revealed a complicated history, including loss in some lineages, movement between chromosomes, and fusion with a non-protein-coding gene. Studying both the functions and evolutionary histories of new proteins illustrates how they might evolve critical roles in biological processes despite their relative novelty. Furthermore, the study of *atlas* identifies an essential genetic player in the fly testis, an important model system for understanding how gametes are produced.

## Introduction

The evolution of new genes is integral to the extensive genotypic and phenotypic diversity observed across species. The best-characterized mechanism of novel gene emergence is gene duplication [1, 2]; however, rapid expansion in high-quality genomic resources has provided mounting evidence of lineage-specific sequences and the existence of alternative modes of new gene origination. One such mechanism is *de novo* evolution, the birth of new genes from previously non-genic or intronic regions, which is now a widely acknowledged source of protein-coding and RNA genes [3–5]. Although *de novo* origination was once considered an unlikely event, catalogs of *de novo* genes have now been published for an expansive range of species [6–13]. Multiple models explain how protein-coding *de novo* genes may acquire both an open reading frame (ORF) and regulatory sequences permitting transcription [14–17]. Interrogation of the biochemical and biophysical properties of the proteins encoded by *de novo* genes has offered initial insight into the mechanisms of emergence and functional potential of these genes [17–20].

The capacity of protein-coding *de novo* genes to evolve important functions is a topic of interest from evolutionary, physiological and molecular perspectives [21]. In the last couple of decades, the products of *de novo* genes have been shown to play diverse roles in a variety of organisms. For example, *de novo* genes function in fundamental molecular processes in yeast, such as *BSC4*, a gene implicated in DNA repair, and *MDF1,* which mediates crosstalk between reproduction and growth [22, 23]. *De novo* genes also evolve roles in organismal responses to disease and changing environmental factors. A putatively *de novo* evolved gene in rice regulates the plant’s pathogen resistance response to strains causing bacterial blight [24]. Antifreeze glycoprotein genes, essential for survival in frigid ocean temperatures, evolved *de novo* in the ancestor of Arctic codfishes to coincide with cooling oceans in the Northern Hemisphere [25, 26]. *De novo* genes are additionally implicated in the development and physiology of mammals. In house mice, a *de novo* evolved gene expressed in the oviduct functions in female fertility by regulating pregnancy cycles [27]. A *de novo* gene found in humans and chimpanzees regulates the oncogenesis and growth of neuroblastoma, revealing the relevance of novel genes to human disease [28]. These studies have started to demonstrate the significance of *de novo* genes, thereby challenging previous assumptions that only ancient, highly conserved genes can be essential.

Across multicellular animals, male reproductive tissues serve as hubs for new gene emergence via numerous mechanisms, including *de novo* evolution [12, 29–34]. Proposed causes of this “out of the testis” phenomenon include the high level of promiscuous transcription in testis cells [35, 36], the relative simplicity of promoter regions driving expression in the testis [37], and preferential retention of novel genes with male-biased expression [38]. Sexual selection also drives rapid evolution of reproductive proteins [39] and could drive the emergence of new genes as a mechanism of improving male reproductive ability [33]. The testis-biased expression of novel genes, combined with growing evidence for new genes acting across a variety of tissue contexts, suggests that many novel genes may function in male reproduction. For example, a pair of young duplicate genes in *Drosophila, apollo* and *artemis,* are essential for male and female fertility, respectively [40]. Continued efforts to identify and characterize testis-expressed novel genes are consequently critical for understanding the genetic basis of male reproductive phenotypes.

*Drosophila* serves as an ideal system for interrogating the prevalence, sequence attributes, expression patterns, and functions of testis-expressed *de novo* genes. The availability of well-annotated genomes for numerous *Drosophila* species, the tractability of flies to molecular genetics techniques, and our thorough understanding of *Drosophila* reproductive processes facilitate comprehensive analyses of novel fly reproductive proteins [41, 42]. As observed in other biological systems, *Drosophila de novo* genes retained by selection demonstrate enriched expression in the testis [10, 19, 33, 34]. The expression patterns of emerging *de novo* genes in the *Drosophila* testis were recently analyzed at single cell resolution [43], thereby providing insight into the dynamics of novel gene expression throughout spermatogenesis. In addition to bioinformatic screens that have started to identify *de novo* genes and large-scale expression analyses of testis-expressed genes, RNAi [44] and CRISPR/Cas9-based [45] functional screens have identified putative, testis-expressed *de novo* genes required for fertility. However, a need remains for in-depth experimental and evolutionary characterization of the genes identified in such screens. Detailed examination of the function of *de novo* proteins will enable us to understand how these proteins might integrate into existing gene networks and become essential.

We previously conducted a pilot functional screen of *de novo* genes with testis-enriched expression in *D. melanogaster* and identified two novel genes, *goddard* and *saturn*, that are required for full fertility [46]. *Goddard* knockdown males failed to produce any sperm. *Saturn* knockdown males produced fewer sperm, which were inefficient at migrating to female sperm storage organs. Subsequent characterization of Goddard using null deletion alleles and a biochemically tagged rescue construct showed that the protein localizes to elongating sperm axonemes and that, in its absence, individualization complexes associate less efficiently with spermatid nuclei and do not successfully progress along sperm tails [20]. These data suggested that putative *de novo* genes can evolve essential roles in a rapidly evolving reproductive process, spermatogenesis.

Here, we expanded this functional screen by evaluating whether any of 42 putative *de novo* genes that show testis-enriched expression in *D. melanogaster* are required for male fertility. We identified one gene, which we named *atlas,* whose knockdown or knockout results in nearly complete male sterility. We found that *atlas* encodes a transition protein that facilitates spermatid chromatin condensation. The *atlas* gene in *D. melanogaster* arose when a likely *de novo* evolved protein-coding sequence moved off of the X chromosome and was inserted upstream of a well-conserved non-coding RNA. While the *atlas* protein-coding sequence has undergone multiple, independent gene loss events since its apparent origin at the base of the *Drosophila* genus, the gene has evolved a critical function in *D. melanogaster*. These results underscore the importance of detailed functional and evolutionary characterization in understanding the origins of new protein-coding genes and the selective forces that affect their subsequent evolution.

## Results

### An RNAi screen identifies a putative de novo gene essential for Drosophila male fertility

A previous pilot screen of 11 putative *de novo* evolved, testis-expressed genes identified two genes that are critical for male fertility in *Drosophila melanogaster* [46]. This result, and other recent work [e.g., 40, 47, 48], suggested that lineage-specific, newly evolved genes can rapidly become important for fertility, perhaps by gaining interactions with existing protein networks. To determine more comprehensively the frequency with which potential *de novo* evolved genes become essential for fertility, we identified *de novo* or putative *de novo* evolved genes with testis-biased expression. A previous computational analysis identified genes that are detectable only within the *Drosophila* genus, lack identifiable protein domains, and show no homology to other known proteins through BLASTP and TBLASTN searches [19]. We filtered these genes to identify those expressed exclusively or predominantly in the testis, a common site of *de novo* gene expression in animal species [27, 34, 38, 49]. This resulted in a set of 96 target genes.

We used testis-specific RNA interference to screen these genes for roles in male fertility. We obtained RNAi lines from the Vienna *Drosophila* Resource Center (VDRC) and the Transgenic RNAi Project (TRiP) and constructed additional lines using the TRiP-style pValium20 vector [50], which induces efficient knockdown in the male germline. We tested an RNAi line for each of 57 genes by using the *Bam*-GAL4 driver, which is expressed in the male germline and which we enhanced with a copy of UAS-*Dicer2*. RT-PCR confirmed at least partial knockdown in lines representing 42 genes (see example in Fig. S1). We then screened knockdown males for fertility by allowing groups of 7 knockdown males to mate with 5 wild-type females for 2 days. Progeny counts were standardized to the number of progeny produced by concurrently mated groups of 7 control males and 5 wild-type females. The results are shown in Fig. 1A. This initial screen identified *CG13541*, whose knockdown severely reduced male fertility. We confirmed the result for *CG13541* by performing single-pair mating fertility assays (Fig. 1B). Consistent with our previous convention of naming testis-expressed genes after American rocketry [46], we will from here on refer to *CG13541* as *atlas*. While RNAi transgenes designed to knockdown *CG43072* and *CG33284* caused full and consistently partial sterility, respectively, we do not further consider these genes because subsequent gene knockout using CRISPR genome editing indicated that neither gene plays a role in male fertility. In these cases, the RNAi phenotypes might have been due to off-target knockdown.

**Figure 1.**
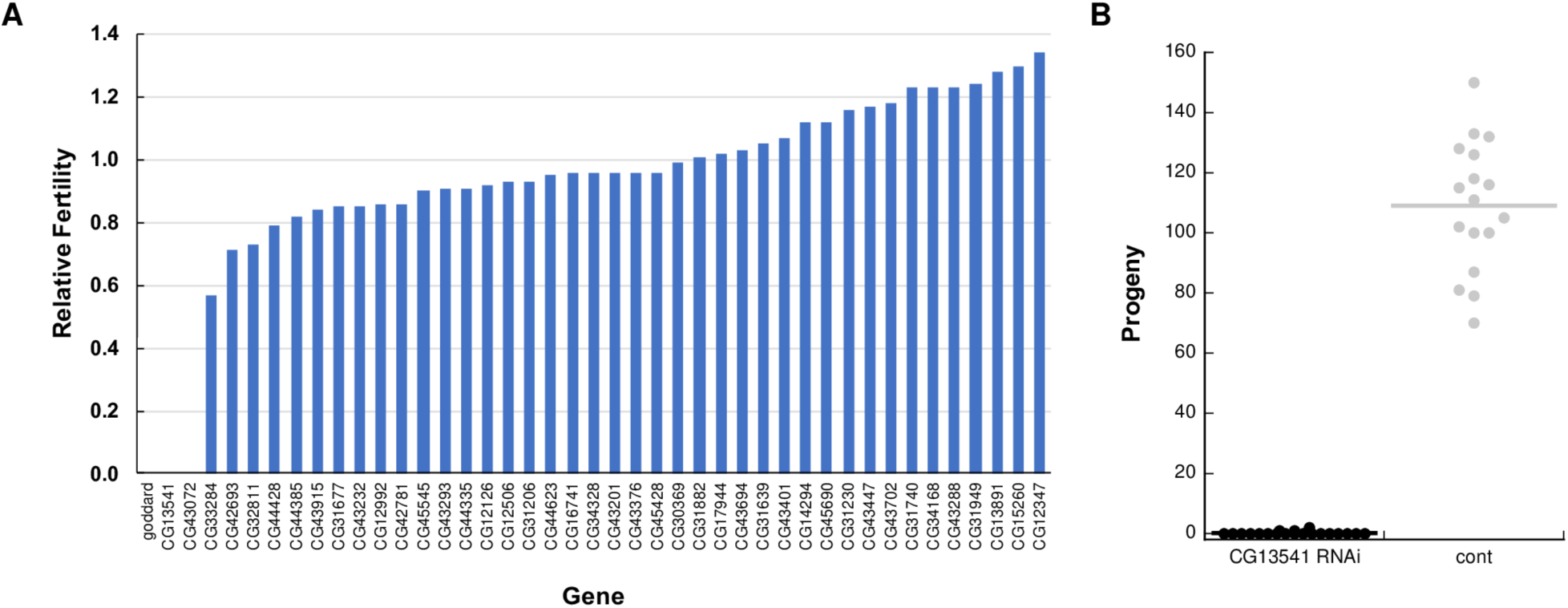
An RNAi screen of putative *de novo* genes identifies *CG13541* as a major contributor to *Drosophila melanogaster* male fertility. A) All RNAi lines that showed at least partial knockdown of the target gene were screened in group fertility assays (see Materials and Methods). Relative fertility was calculated by dividing the average number of progeny produced per female mated to knockdown males by the average number of progeny produced per female mated to control males in a contemporaneous experiment. Relative fertility measurements lack error bars because each gene was tested in only 1-2 replicates. Knockdown of *goddard* was used as a positive control. B) A single-mating, single-pair fertility assay confirms the observed defect when males are knocked down for *CG13541*, as knockdown males showed significantly reduced fertility (control fertility (mean ± SEM): 109.0 ± 5.3; knockdown fertility: 0.2 ± 0.1; two-sample *t*-test assuming unequal variances, *p* = 5.6 x 10^-13^).

### CRISPR-mediated gene mutation validates atlas RNAi results

We validated the observed fertility defect by using CRISPR/Cas9-based genome editing to construct putative loss-of-function alleles for *atlas* (Fig. S2). The principal allele we used for validation and the functional studies described below was a null allele that completely deletes the *atlas* genomic locus. This allele was generated by targeting each end of the locus with a gRNA. We made three additional frameshift alleles by inducing double-stranded breaks at a gRNA target site just downstream of the *atlas* start codon, which induced non-homologous end joining. Males homozygous for the *atlas* deletion allele have the same fertility defect as knockdown males (Fig. 2A). Males homozygous for any of three frameshift alleles showed significantly reduced, but non-zero, fertility (Fig. 2B). It is possible that residual *atlas* function may be present in these animals, perhaps due to translation initiation at a downstream start codon, which could generate a shorter protein with partial function. Each frameshift allele retains the possibility of encoding an N-terminally truncated, but otherwise in-frame, version of Atlas protein that would lack the first 60 amino acids (out of 172; see Fig. S2). Alternatively, it is possible that the residual fertility of the frameshift alleles is caused by the gene’s intact 3’ UTR, a topic we discuss in more detail below. Finally, we constructed a genomic rescue construct carrying both the full *atlas* transcribed region and its native regulatory sequences. *atlas* null males that carried a single copy of the rescue construct had fully restored fertility (Fig. S3). Overall, these data demonstrate that *atlas* loss, and not an RNAi or CRISPR off-target, causes nearly complete male sterility.

**Figure 2.**
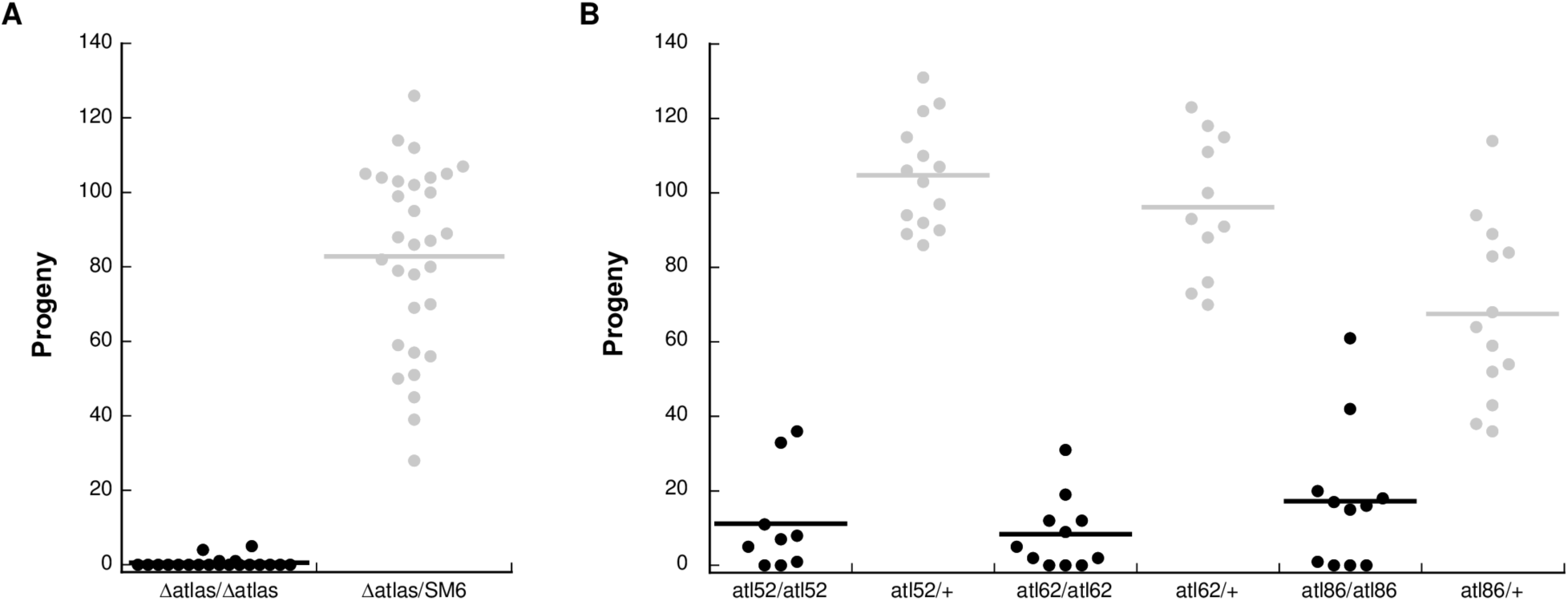
CRISPR-generated deletion and frameshift alleles of *atlas* confirm the gene’s requirement for male fertility. A) Single-pair fertility assay for males homozygous for the null (Δ*atlas*) allele or heterozygous controls (Δ*atlas*/SM6). Flies homozygous for the deletion had significantly reduced fertility (control fertility: 82.9 ± 4.5; null fertility: 0.3 ± 0.6; two-sample t-test assuming unequal variances, *p* = 5.4 x 10^-18^). B) Single-pair fertility assays for males homozygous or heterozygous for three frameshift alleles of *atlas* generated by imprecise non-homologous end joining at a CRISPR/Cas9 target site just downstream of the start codon: *atlas*^52^ (control fertility: 104.7 ± 3.8, mutant fertility: 11.2 ± 4.6; two-sample *t*-test assuming unequal variances: *p* = 8 x 10^-12^), *atlas*^62^ (control fertility: 96.2 ± 5.7; mutant fertility: 8.4 ± 2.9; two-sample *t*-test assuming unequal variances: *p* = 6.1 x 10^-10^) and *atlas*^86^ (control fertility: 67.5 ± 6.6; mutant fertility: 17.3 ± 5.8; two-sample *t*-test assuming unequal variances: *p* = 9.5 x 10^-6^).

### Atlas is required for proper spermatid nuclear condensation

We next examined how *atlas* loss of function impacted male fertility at the cellular level. Dissection and phase-contrast imaging of *atlas* deletion null or knockdown male reproductive tracts revealed that while the pre-meiotic and meiotic stages of spermatogenesis appeared normal, sperm accumulated at the basal end of the testes, rather than in the seminal vesicles (SVs), over the first week of adulthood (Fig. 3A and Fig. S4). To further characterize the fertility defects in the absence of *atlas*, we examined the Mst35Bb-GFP (“protamine”-GFP) marker in null or knockdown backgrounds [51]. Mst35Bb encodes one of two protamine-like proteins (highly similar paralogs of each other) that bind DNA in mature sperm. Its GFP fusion construct thus allows visualization of nuclei in late stage spermiogenesis and mature sperm. Consistent with the observed conglomeration of sperm tails at the basal testes, SVs from either *atlas* null or knockdown males contained fewer mature sperm (Fig. 3B, Fig. S4C). The nuclei of sperm from null males also appeared wider and less elongated than those of controls. Together, these data suggest that *atlas* is required after meiosis, as developing spermatids take on their final structures.

**Figure 3.**
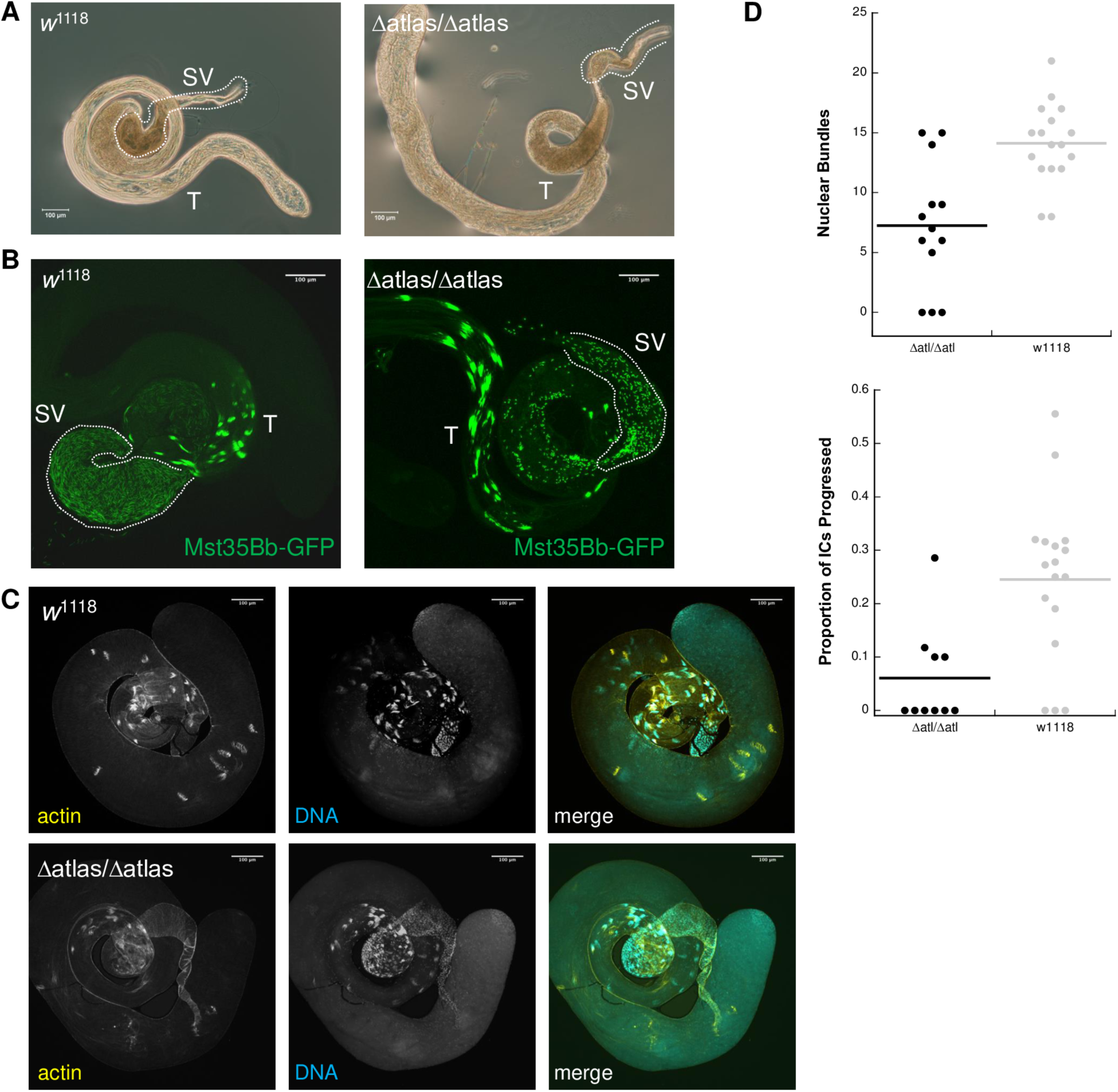
Cytological investigations of the *atlas* mutant fertility defect. A) Phase contrast microscopy of male reproductive tracts dissected from 7-day-old, unmated control (*w*^1118^) or *atlas* null males. Control males show the expected accumulation of sperm tails in the seminal vesicle (SV), which appear here as a darker brown shading, while null male have an aberrant accumulation of sperm tails at the basal end of the testis (T). B) Visualization of Mst35Bb-GFP in 4-day old control and *atlas* null testes. While Mst35Bb localizes to spermatid nuclei in the absence of *atlas*, the nuclei appear shorter and much less numerous in the outlined SV. C) Representative images from phalloidin staining of *w*^1118^ and *atlas* null testes used to assess the association of individualization complexes (ICs) with nuclear bundles and the progression of ICs down the length of sperm tails. D) At top, number of nuclear bundles with ICs associated in control and *atlas* null testes. Significantly more ICs were observed in control testes (control: *N* = 17, median = 14; mutant: *N* = 13, median = 7; Wilcoxon rank-sum test W = 34, *p* = 0.0014). At bottom, proportion of all observed ICs that were intact and that had progressed away from nuclear bundles. Three mutant testes with no observed ICs were excluded from the analysis. A significantly higher proportion of ICs progressed in control testes (control: *N* = 17, median = 0.27; control: *N* = 10, median = 0; Wilcoxon rank-sum test W = 28, *p* = 0.0038).

We next examined two post-meiotic processes: individualization of 64-cell spermatid cysts into mature sperm, and spermatid nuclear condensation. Individualization initiates when an actin-rich individualization complex (IC) associates with the bundle of spermatid nuclei. The IC then proceeds down the sperm tails, expelling cytoplasmic waste and remodeling cell membranes to form 64 individual sperm. We visualized this process in males 0-1 days old, when spermatogenesis occurs at high levels, by staining whole mount testes for actin (Fig. 3C-D). Although ICs associated with nuclear bundles present at the basal end of the testes in both control and *atlas* null males, we observed significantly fewer nuclear bundle-associated ICs in nulls (Fig. 3C-D). While control testes typically had several ICs progressing down sperm tails, we saw a significantly reduced proportion of progressed bundles in nulls (Fig. 3C-D). In some null testes, we also observed individual investment cones dissociated from progressing ICs (Fig. 3C).

The ability of ICs to assemble at nuclear bundles and progress down sperm tails may be reduced if nuclear condensation is aberrant [reviewed in ref. 52]. During *Drosophila* spermiogenesis, round spermatid nuclei undergo a series of stepwise, morphological changes that are the product of two distinct, but related processes: changes in the chromatin packaging of DNA, and changes in nuclear shape [53–55]. The end result is thin, condensed nuclei. We quantified this process in testes dissected from newly eclosed wild-type and *atlas* null males expressing Mst35Bb-GFP, which marks the final stages of condensation. We shredded the post-meiotic region of the testes in the presence of a fixative and counted the number of nuclear bundles that exhibited each of five stages of condensation [53]: round nuclei, early canoe-stage (unmarked with Mst35Bb-GFP), late canoe-stage (marked with Mst35Bb-GFP), elongated nuclei, and fully condensed nuclei (Fig. 4). Condensation of the nuclear bundles in *atlas* null testes progressed at similar rates to controls through the late canoe stage (Table S1). However, in *atlas* null males, all nuclear bundles that progressed past the canoe stage (which included ∼60% [range: 26-100%] of all observed bundles) showed an aberrant “curled” phenotype (Fig. 4; Table S1). These data suggest that Atlas protein is required during the later stages of nuclear condensation and are consistent with the idea that the loss of *atlas* affects nuclear condensation in a way that reduces IC assembly and sperm individualization (see Fig. 3C).

**Figure 4.**
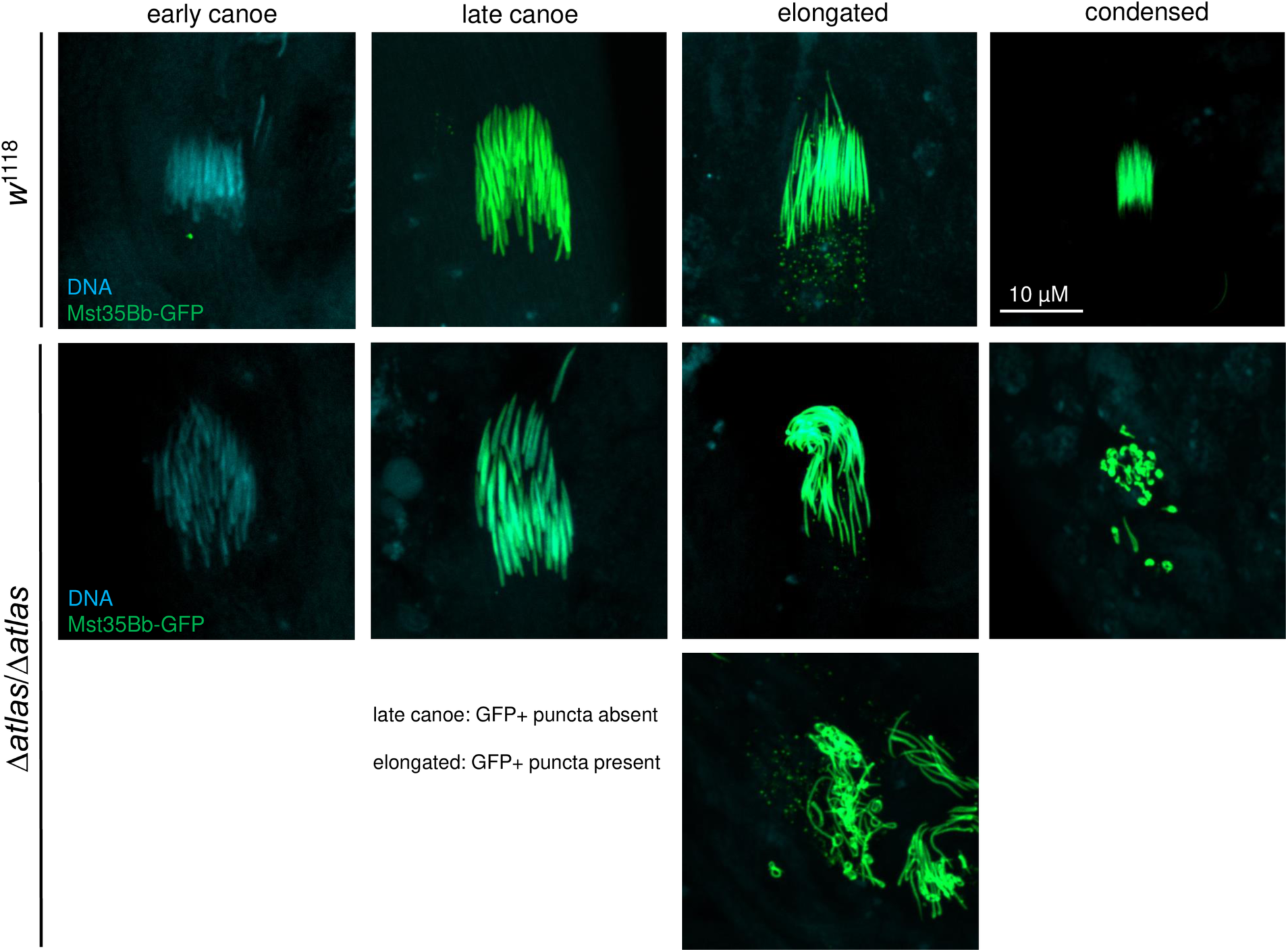
*Atlas* null males show aberrant nuclear shaping at and beyond the elongated stage of spermatid nuclear condensation. Early and late canoe stages were distinguished by the absence or presence of Mst35Bb-GFP, respectively. Late canoe and elongated stages were distinguished by the absence or presence of GFP-positive puncta, respectively. Condensed nuclei were distinguished from elongated nuclei by size. As shown in Table S1, nuclear bundles from *atlas* null testes consistently took on a curved shape after the canoe stage, though the degree of curvature was variable, as exemplified by the two examples of elongated nuclei from *atlas* null testes above.

That condensing spermatid nuclei are misshapen in the absence of *atlas* suggests the possibility that Atlas protein is critical for nuclear condensation. This idea is further supported by its predicted biochemical properties. Previously characterized spermatid chromatin binding proteins are small and highly basic [53, 56, 57], as the excess of positively charged amino acid side chains facilitates ionic interactions with negatively charged DNA. Many such proteins (i.e., Tpl94D, Mst35Ba, Mst35Bb, Prtl99C and Mst77F) also contain a conserved protein domain, the high-mobility-group box (HMG-box) domain [55, 58–64], suggesting that this type of chromatin binding protein could have originated through gene duplication and divergence. Consistent with its putative *de novo* origin, Atlas lacks a detectable HMG-box domain. However, Atlas is otherwise similar to these other sperm chromatin binding proteins: the ∼20 kDa protein has a highly basic predicted isoelectric point of 10.7, and its primary sequence contains the sequence KRDK, which matches the canonical consensus sequence for nuclear import, K(K/R)X(K/R) [65]. To test the hypothesis that Atlas is nuclear localized, and could thus bind DNA, we generated an *atlas-GFP* transgene under UAS control and expressed it ubiquitously using *tubulin*-GAL4 and in the early male germline using *Bam*-GAL4. In both larval salivary glands and early male germline cells, Atlas-GFP appeared to be nuclear localized (Fig. S5).

While these results were consistent with Atlas protein localizing to the nucleus, they did not allow us to visualize Atlas in the cells in which it is normally expressed. To do so, we used CRISPR/Cas9-induced homology directed repair (https://flycrispr.org/scarless-gene-editing/) [66–68] to create an *atlas*-GFP fusion at the endogenous *atlas* locus (see Fig. S6 and Materials and Methods). We first confirmed the functionality of the knock-in allele by showing that males with the *atlas* locus genotype *atlas-*GFP/Δ*atlas* had equivalent fertility to males of genotype *atlas+*/Δ*atlas* (Fig. 5A). We then visualized Atlas-GFP fusion protein in whole-mount testes in conjunction with phalloidin-stained actin (Fig. 5C and Fig. S7A). Atlas-GFP was absent from seminal vesicles, consistent with its absence from the proteome of mature *D. melanogaster* sperm [69, 70]. Instead, Atlas-GFP colocalized with condensing nuclear bundles near the basal end of the testes (Fig. 5C). Actin-based ICs were also observed in the basal testes, but generally did not co-localize with Atlas-GFP, suggesting that Atlas-GFP is present in condensing nuclei before IC association (Fig. 5C). This result, taken together with the aberrant nuclear condensation in the absence of *atlas* (Fig. 4), is consistent with the idea that Atlas is a transition protein. Transition proteins are chromatin components that act transiently during spermatid nuclear condensation. A series of transition proteins first replace histones as the primary DNA binding proteins in the nucleus and then give way to protamines, the proteins that package chromatin in mature sperm [53, 55, 58].

**Figure 5.**
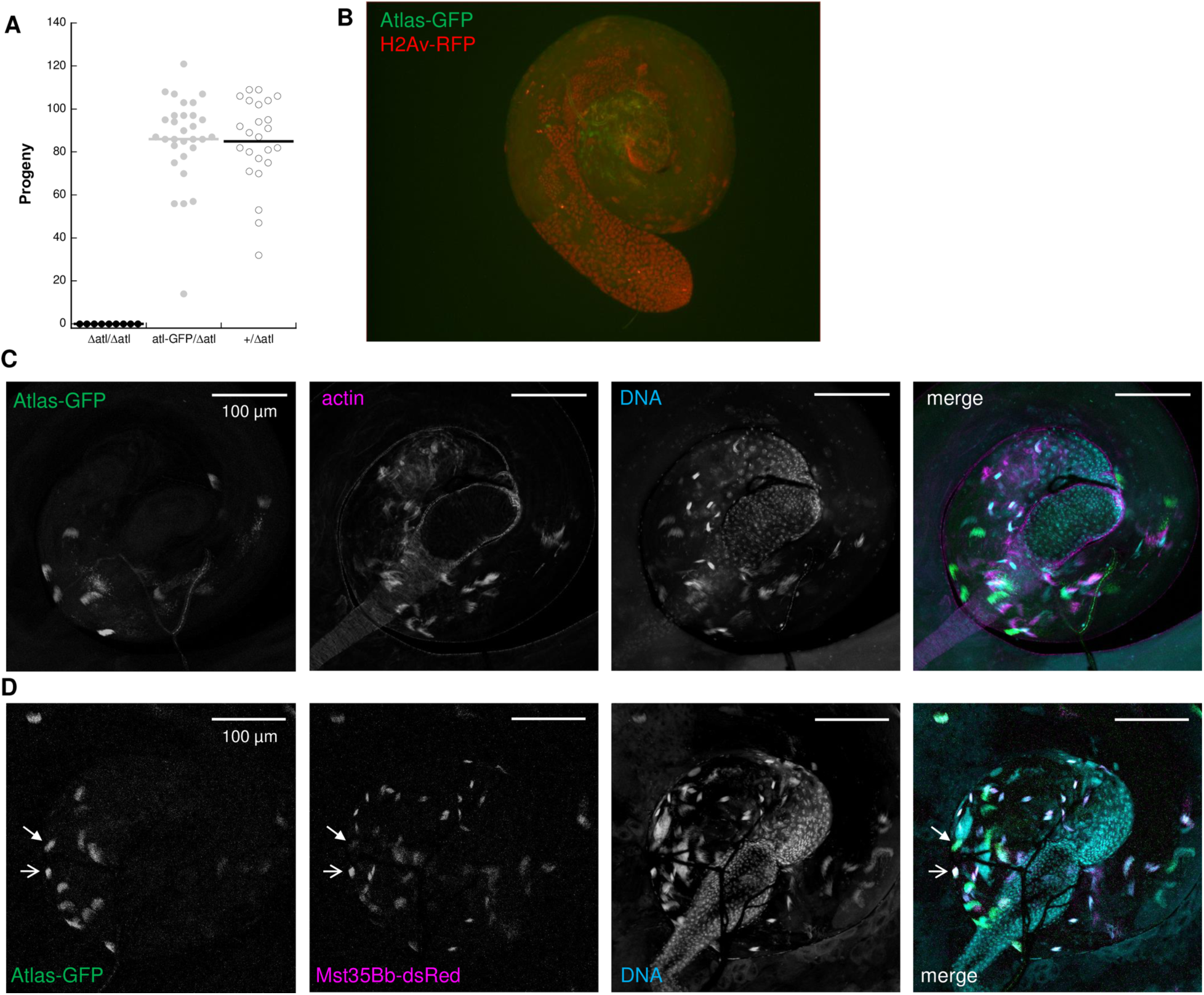
An *atlas*-GFP allele generated at the endogenous *atlas* locus is fully functional for male fertility and encodes a protein that localizes to condensing spermatid nuclei. A) A single copy of the *atlas-*GFP allele is sufficient for full fertility when paired with the Δ*atlas* allele, as compared to males heterozygous for the wild-type *atlas* allele (Δ*atlas*/*atlas*-GFP fertility: 86.0 ± 3.2; Δ*atlas*/+ fertility: 84.9 ± 4.1; two-sample t-test assuming unequal variances, *p* = 0.85). B) Atlas-GFP does not co-localize with histone H2Av-RFP, a marker of the initial stages of spermatid nuclear condensation. C) Visualization of Atlas-GFP in the basal portion of whole-mount testes from *atlas-GFP* homozygotes shows that the fusion protein co-localizes with a subset of condensing spermatid nuclear bundles. While actin associates with fully condensed nuclei at the basal testis, Atlas-GFP does not overlap and is also absent from the seminal vesicle. D) Atlas-GFP partially colocalizes with Mst35Bb-dsRed, a marker of the final stage of nuclear condensation, in the basal portion of whole-mount testes. Open arrow: example of co-localization. Filled arrowhead: example of Atlas-GFP that does not co-localize with Mst35Bb-dsRed. Collectively, these data suggest that atlas may serve as a transition protein involved in the final stages of nuclear condensation. The whole testes from which the basal portions are shown in panels C and D are shown in Fig. S7.

To further elucidate the role of *atlas* in nuclear condensation, we next examined Atlas-GFP localization in the presence of either an early spermatid nuclear marker, histone H2Av-RFP [55, 71], or Mst35Bb-dsRed [51], a marker of nuclei from the late canoe stage through final condensation. Atlas-GFP showed no co-localization with H2Av-RFP, suggesting that Atlas functions after histone removal (Fig. 5B). In contrast, some GFP-positive bundles co-localized with Mst35Bb-dsRed, but others did not (Fig. 5D and Fig. S7B). These data suggest that Atlas may be one of the final transition proteins used in nuclear condensation before the chromatin becomes fully condensed with protamines.

To determine the stage(s) of nuclear condensation at which *atlas* functions, we analyzed the shape of fixed nuclear bundles from shredded testes isolated from *atlas*-GFP males on the day of eclosion. Based on the stage of the defect in *atlas* null males (Fig. 3-4) and the pattern of Atlas-GFP-positive bundles in whole-mount testes (Fig. 5), we hypothesized that Atlas-GFP would localize to the later stages of nuclear condensation. Consistent with this hypothesis, we did not detect Atlas-GFP in round or early canoe stage bundles (Fig. 6A-B). Atlas-GFP co-localized with DNA in late canoe stage bundles (Fig. 6C). Interestingly, when nuclei elongated further, GFP was detected not in the nucleus, but as puncta basal to the nuclei (Fig. 6D; see also Fig. S7A). Since Atlas-GFP is not observed in mature sperm in the SV (Fig. 5C), these data suggest that Atlas may function as a transition protein that facilitates the condensation of spermatid nuclei from histone-based DNA packaging to protamine-like-based DNA packaging [53] and is then removed from nuclei once protamines bind DNA. Indeed, the appearance of Atlas in nuclei during the late canoe stage of condensation is similar to the pattern observed for a previously characterized transition protein, Tpl94D [53]. We hypothesize that the failure of *atlas* null sperm to form needle-like nuclei can be explained by the absence of Atlas from the late canoe nucleus. It is also possible that the apparent removal of Atlas-GFP from nuclei (Fig. 6D) represents a mechanism for removing transition proteins from the nucleus after they exert their functions. We observed above that some Mst35Bb-GFP also appears to be removed from the nucleus in puncta during the elongation stage of nuclear condensation (see elongated stage of control nuclear bundles, Fig. 4), even though other Mst35Bb-GFP molecules ultimately package DNA in mature, individualized sperm. This could occur if Mst35Bb-GFP is present in excess of what is needed to package DNA.

**Figure 6.**
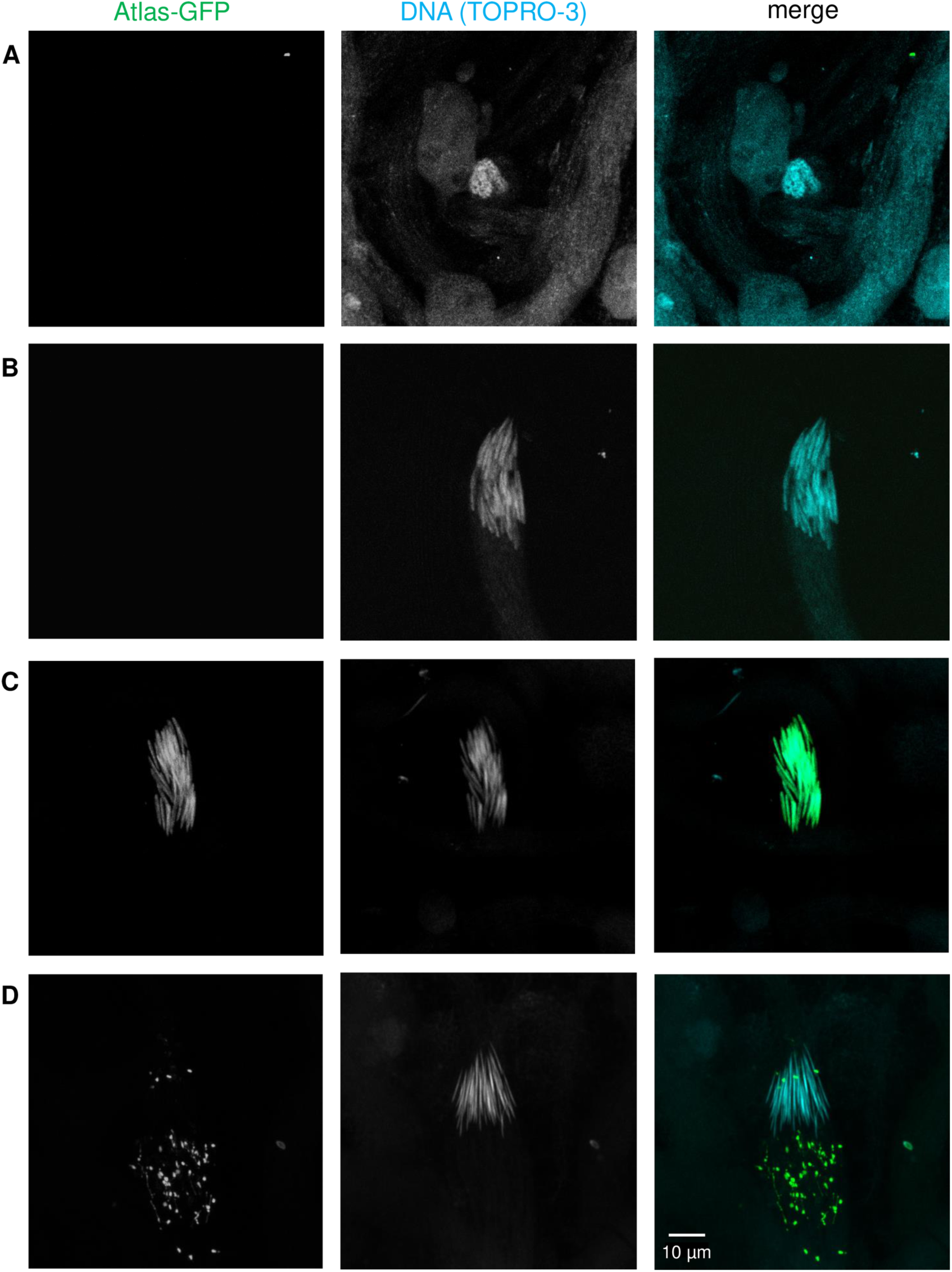
Atlas-GFP is present in late canoe-stage spermatid nuclei and then appears to leave the nucleus in puncta. Staging of condensing spermatid nuclei fixed in paraformaldehyde from *atlas*-GFP males stained with TO-PRO-3 DNA stain. Atlas-GFP is not detectable in (A) round stage or (B) early canoe stage nuclei. Atlas-GFP is nuclear localized in the late canoe stage (C). When nuclei become fully elongated (D), puncta of Atlas-GFP appear to be removed from the nucleus.

### Evolutionary origins of atlas

To better understand the evolutionary origin of *atlas* and its evolution since emergence, we used a combination of BLAST- and synteny-based approaches to identify *atlas* orthologs throughout the genus [46, 72]. One notable feature of this two-exon gene is that the protein-coding region (519 nucleotides) is contained entirely within the first exon (622 nt); the longer, second exon (910 nt) appears to be entirely non-coding (Fig. 7). Surprisingly, the second exon is more widely conserved. BLASTN detected significant matches to this region (range of hit length: 185-864 nt) in the same genomic location on Muller element C [see ref. 73 for explanation of Muller elements], as assessed by synteny, in all *Drosophila* species examined, including distantly related species such as *D. virilis* and *D. grimshawi* (Fig. 7 and S8). The protein-coding first exon shows a more limited phylogenetic distribution. In most members of the *melanogaster* group of *Drosophila* (gray box in Fig. 7), this exon is found in a conserved position, adjacent to the non-coding region on the equivalent of *D. melanogaster* chromosome 2R (Table S2). In *D. ananassae*, however, the protein-coding region is found on the X chromosome (Muller element A). A putative ortholog for the protein-coding sequence is detectable by BLASTP in *D. virilis* in a partially syntenic region on the same Muller A element (Table S2, Fig. S9). These data suggest that the *atlas* protein-coding sequence initially arose on Muller element A and then moved to Muller element C, giving rise to the gene structure observed in extant *D. melanogaster* and its sister species.

**Figure 7.**
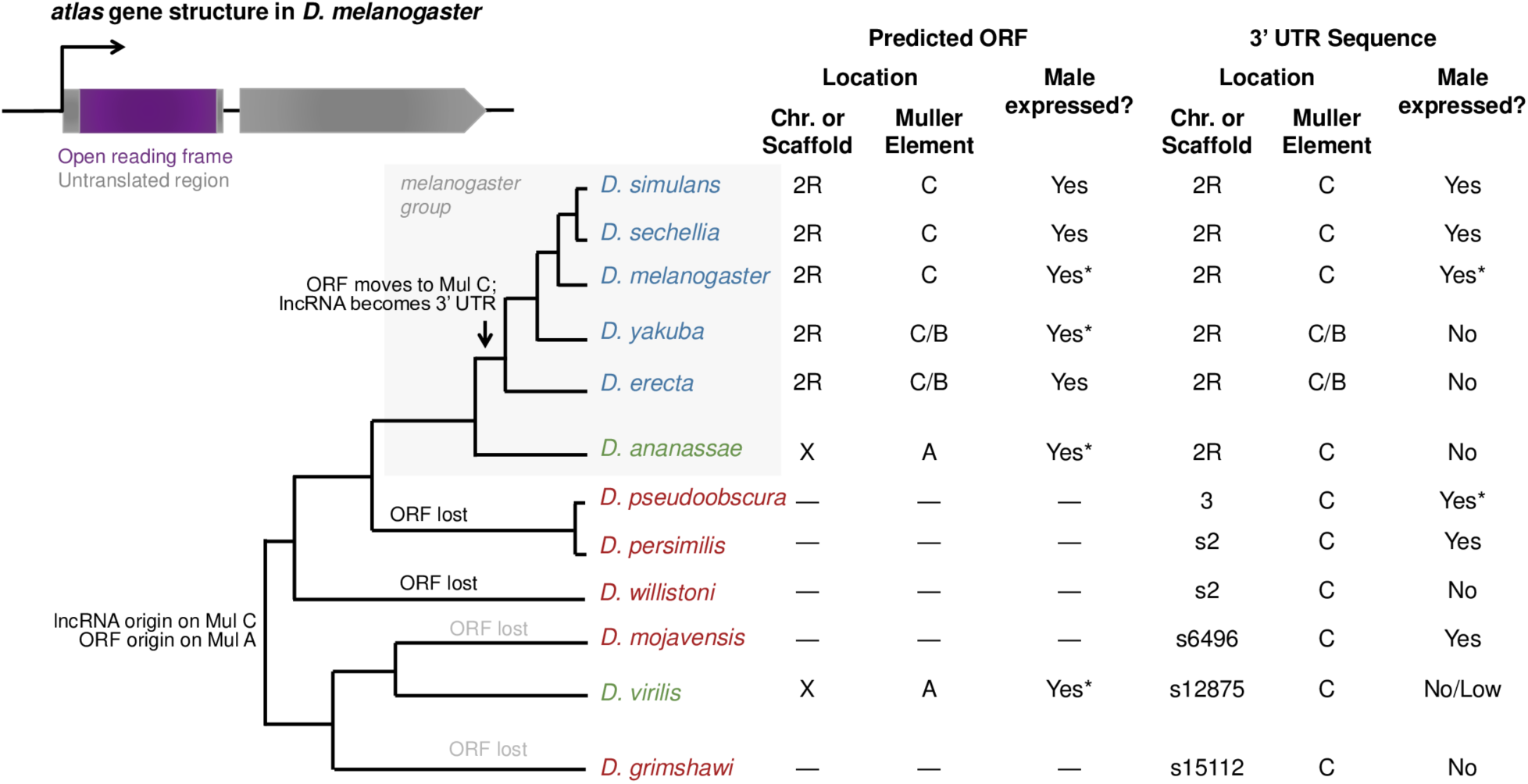
Molecular evolution and gene expression of *atlas* across the *Drosophila* genus. The gene structure of *atlas* in *D. melanogaster* is shown at top left. The predicted protein-coding sequence is contained entirely within exon 1, while exon 2 encodes the presumed 3’ UTR. The gene is located on chromosome 2R, equivalent to Muller element C. The phylogeny shows BLAST- and synteny-based detection of sequences orthologous to the protein-coding sequence and the 3’ UTR sequence across *Drosophila* species. Sex-specific adult RNA-seq data were used to assess male expression across species, with RT-PCR verification performed in species marked with asterisks. RNA-seq data for the syntenic region of the 3’ UTR in *D. virilis* were ambiguous; see Figs. S8 and S10.

To confirm the lack of *atlas* protein-coding sequences identifiable by BLASTP or TBLASTN in most non-*melanogaster* group species, we identified the regions syntenic to those containing *atlas* in *D. ananassae* and *D. virilis* in 11 additional *Drosophila* species and used more sensitive methods to search for potential orthologs [72]. Specifically, we: a) relaxed the BLAST cut-offs for detection, since default parameters can cause false-negative results when searching for potential *de novo* genes in divergent species [74]; b) used adult male RNA-seq data to detect transcribed areas within each syntenic region that did not match annotated genes; and, c) predicted the isoelectric point of the potential proteins encoded, under the hypothesis that Atlas orthologs would have conserved, DNA-binding functions. The results are summarized in Fig. S9 and Table S2. These searches detected no evidence for *atlas* orthologs in the following *Drosophila* species: *obscura*, *miranda, willistoni, hydei, arizonae, mojavensis, navojoa,* and *grimshawi*. Some of these species have unannotated, male-expressed transcripts in the regions syntenic to the Muller A location of *atlas* in *D. ananassae* and *D. virilis*, but when each was compared with BLASTP to the *D. melanogaster* proteome, all matched proteins other than Atlas, suggesting they may be lineage-specific paralogs of other genes (Fig. S9). In sister species *D. pseudoobscura* and *D. persimilis*, we detected a male-expressed transcript predicted to encode a protein with a pI > 10 in the region syntenic to the location of *atlas* in *D. virilis*, but the predicted protein sequences showed no significant BLASTP similarity to *atlas* orthologs (Table S2). While this predicted protein may represent a divergent *atlas* ortholog, the abSENSE method predicts low probabilities of BLASTP detection failure when searching for Atlas protein in these species (0.02 and 0.04, respectively), so we favor the hypothesis of a lineage-specific, newly evolved gene in the region. Conversely, in *D. busckii*, we detected, in the region syntenic to the *D. virilis atlas* locus, a male-expressed transcript predicted to encode a protein with significant BLASTP identity to *D. melanogaster* Atlas (e = 4e-8), but with a predicted pI of 5.1 and a ∼50 percent shorter open-reading frame (Table S2). The ortholog status of this predicted protein is also unclear, but because of its dramatically altered size and pI, it is unlikely to have a functional role equivalent to that of *D. melanogaster* Atlas.

To investigate whether the protein-coding region may have reproductive functions in other species, we used sex-specific RNA-seq data from numerous *Drosophila* species curated by the Genomics Education Partnership [72; thegep.org] and verified several of these results by RT-PCR (Fig. 7, Fig. S10). In all species in which *atlas* was detected, the protein-coding region is expressed specifically in males regardless of its genomic location (Fig. 7). Interestingly, the non-coding region shows male-specific expression in species lacking an unambiguous, orthologous coding region, such as *D. pseudoobscura* and *D. mojavensis*. Conversely, while *D. yakuba* and *D. erecta* express the protein-coding region robustly, we found no RNA-seq evidence to support expression of the non-coding second exon, in spite of its sequence conservation (Fig. S8). Based on its high level of sequence conservation, consistent genomic location and expression in a variety of species, it is possible that what we now consider to be the 3’ untranslated region of *atlas* from *D. melanogaster* was, ancestrally, a non-coding RNA.

The FlyBase database reports two transcript isoforms of *atlas* in *D. melanogaster*: the *atlas*-RA isoform is 986 nucleotides, while the *atlas*-RB isoform is 1528 nt. These isoforms differ in how much of the second, non-coding exon is included in the transcript. We used RT-PCR of whole male cDNA to assess the presence of these isoforms and their relative abundances. Primers designed to amplify a region present in both isoforms produced products that appeared more abundant than primers designed to amplify only the long isoform, even though both primer pairs appeared to amplify genomic DNA with equal efficiency. Based on RT-PCR band intensities and controlling for product size and genomic PCR band intensities, we estimated that the short isoform is about 3-fold more abundant. This difference in abundance is mirrored in available RNA-seq data, which show approximately 3- to 4-fold higher levels of expression in the upstream part of exon 2 (Fig. S8), a pattern that also appears in *D. simulans* and *D. sechellia*. Evaluating the potential significance of this finding awaits functional characterization of the non-coding region.

As we have observed for other putative *de novo* genes with essential male reproductive functions [46], the pattern of *atlas* protein-coding sequence presence/absence across the phylogeny is difficult to explain parsimoniously. If we assume that gene birth events are less frequent than gene deaths, since the latter can occur through many possible mutational events and can happen separately along multiple phylogenetic lineages, our data support the hypothesis of a single origin of the protein-coding sequence at the base of the genus, followed by independent loss events on the lineages leading to *D. grimshawi*, *D. mojavensis* and *D. willistoni*, and potentially also *D. pseudoobscura*/*persimilis* (Table S2). We summarize these findings for 12 representative species of *Drosophila* in Fig. 7. The general patterns of loss do not change when all species of Table S2 are considered, though an additional loss in the *melanogaster* group is likely due to the absence of a detectable ortholog in *D. kikkiwai* and *D. serrata*. As noted above, the pattern of gene loss can also appear due to orthology detection failure [74], for which we tried to account with our additional search methods described above. We also note, however, that the probability of BLASTP-based ortholog detection failure is relatively low for some *Drosophila* species that lack *atlas*, including *D. pseudoobscura* (probability of non-detection due to divergence = 0.02), *D. persimilis* (p = 0.04) and *D. willistoni* (p = 0.06). The probability is higher for other species, *D. mojavensis* (p = 0.33) and *D. grimshawi* (p = 0.66), underscoring the importance of our additional search strategies. Overall, our data support the hypothesis of multiple, independent loss events within *Drosophila*.

AbSENSE produces a 1.00 probability of BLASTP-based Atlas ortholog detection failure outside of *Drosophila*, reflecting the protein’s short length and relatively rapid divergence (see below). Indeed, the protein-coding and non-coding transcriptomes from each species showed no matches to Atlas protein or cDNA sequences by BLAST. We thus used another synteny-based approach, summarized in Fig. S11, to look for the protein-coding gene in other Dipterans with well-resolved genomes: *Musca domestica*, *Glossina morsitans*, *Lucilia cuprina*, *Aedes aegypti*, *Anopheles darlingi*, *Anopheles gambiae*, *Culex quinquefasciatus* and *Mayetiola destructor*. In none of these species was a putative homolog found in any potential syntenic region.

Recognizing the limitation of even this approach, we also used HMMER [75] to search iteratively either all genomes in ENSEMBL, or all metazoan genomes in ENSEMBL, for annotated proteins with identity to Atlas from *D. melanogaster* or *D. virilis*. These searches initially identified significant hits to the Atlas orthologs we identified above from other *Drosophila* species. When these collections of orthologs were used as queries, no further proteins outside of *Drosophila* were a significant match. As a control, we performed the same search strategy with *D. melanogaster* Mst35Bb, a protein whose length, amino acid composition, and function are similar to Atlas. These searches readily identified orthologs throughout Diptera, consistent with predictions of its conservation from the OrthoDB database [76]. We also note that a previous HMMER-based analysis to identify HMG-box-containing spermatid chromatin condensation proteins in *Drosophila* and other related insects did not detect *atlas* as a member of the gene family [64], further diminishing the possibility that *atlas* is a divergent paralog of other transition and protamine-like proteins. Thus, we conclude that *atlas* is a putative *de novo* evolved gene that is limited phylogenetically to the *Drosophila* genus.

Finally, we used standard tests of molecular evolution to examine the selective pressures that have shaped Atlas protein within the *melanogaster* group. We aligned the *atlas* protein-coding sequences from 12 species and used PAML to ask whether a model (M8) allowing for positive selection, as well as neutral evolution and purifying selection, explained the data better than models (M7 and M8a) that allowed only neutral evolution and purifying selection [77, 78]. These data showed that while the *atlas* protein-coding sequence’s rate of evolution was accelerated relative to most *Drosophila* proteins (whole-gene estimated *d*_N_/*d*_S_, ω = 0.41 by PAML model M0), there was no significant evidence for positive selection acting to recurrently diversify a subset of sites within the protein (Table 2).

**Table 2.** PAML sites tests for positive selection acting on atlas in the melanogaster group.

| PAML Model | $\omega$ estimate | ln L | Number of parameters | Likelihood ratio test with M8 |
| --- | --- | --- | --- | --- |
| M0 (uniform $\omega$ across all sites) | 0.41 | -4032.24 | 23 | n/a |
| M7 (10 site classes, each with $0 \leq \omega \leq 1$) | n/a | -3961.28 | 24 | $\chi^2 = 1.74$, df = 2, $p = 0.42$ |
| M8a (10 site classes as in M7, plus one class with $\omega = 1$) | n/a | -3961.02 | 25 | $\chi^2 = 1.22$, df = 1, $p = 0.27$ |
| M8 (10 site classes as in M7, plus one class with $\omega \geq 1$) | 1.39 (9.1% of sites) | -3960.41 | 26 | n/a |

## Discussion

Across taxa, many *de novo* evolved genes are expressed in the male reproductive system [12, 31, 33, 34]. Identifying those genes that have evolved essential roles in reproduction will provide insight into how newly evolved genes integrate with existing cellular networks [47] and how evolutionary novelties permit adaptation in the face of sexual selection. Here, we screened 42 putatively *de novo* evolved genes for major effects on male *D. melanogaster* reproduction. Our primary screen identified three genes whose knockdown caused an apparent reduction in male fertility. However, subsequent CRISPR-mediated gene deletion revealed that only one of these genes, *atlas*, was truly essential. This result underscores the importance of validating genes identified in RNAi screens through traditional loss-of-function genetics and other approaches.

Loss *of atlas* function reduces fertility by affecting mature sperm production. During spermiogenesis, *atlas* mutants show aberrant nuclear condensation and an inability to individualize spermatid bundles successfully. GFP-tagged Atlas protein localizes to condensing spermatid nuclei in the basal testis and partially co-localizes with Mst35Bb, a protamine around which DNA is wrapped in mature, individualized sperm. Evolutionary analysis showed that the *atlas* protein-coding sequence likely arose at the base of the *Drosophila* genus but was unlikely to have played an essential role immediately upon birth, as the gene was subsequently lost along several independent lineages. Within the *melanogaster* group of *Drosophila*, however, the gene moved from the X chromosome to an autosome, where it formed a single transcriptional unit with a conserved, non-coding sequence. Since this point, the gene has encoded a protein with a conserved length, isoelectric point and male-specific expression pattern, suggesting potential functional conversation over the last ∼15 million years.

### Comparison of atlas null and frameshift allele phenotypes

The null deletion and frameshift alleles of *atlas* all caused significantly reduced fertility (Fig. 2), but the deletion allele resulted in essentially complete sterility, while residual fertility remained in males homozygous for each frameshift allele. We noted above and depicted in Fig. S2 how the frameshift alleles have the potential to encode an N-terminally truncated form of Atlas that would contain amino acids 61-172 of the wild-type protein, if these alleles allow translation initiation at the methionine-encoding codon 61. Such a truncated protein would be 35 percent shorter than wild-type, lack the predicted nuclear localization sequence, and have a reduced isoelectric point of 7.1. Each of these factors could contribute to reduced functionality.

The broad sequence conservation of the 3’ UTR across the *Drosophila* genus, including in species that lack the *atlas* coding sequence, suggests the alternative possibility that the 3’ UTR also contributes to fertility. In one scenario, the 3’ UTR could act as part of the *atlas* locus by regulating RNA stability and/or the spatiotemporal control of Atlas protein translation. An alternative possibility is that the 3’ UTR functions on its own as a non-coding RNA. Indeed, since the protein-coding region and the *D. melanogaster* 3’ UTR were initially separate genetic entities (and continue to be in *D. ananassae* and *D. virilis*), it is possible that each contributes to fertility in a unique way. Future experiments that could distinguish these possibilities are described below.

Based on the current evidence, however, we think that the primary way in which *atlas* impacts fertility is through its protein-coding sequence. This conclusion is supported by: the ∼60-90 percent reduction in fertility in even the frameshift mutants; the stability of the Atlas-GFP fusion protein and its presence in the spermatid nuclear condensation stages that immediately precede the timing of the null mutant phenotype; and, the observation that the protein-coding region shows a more highly conserved expression pattern than the UTR region within the *melanogaster* group (Fig. 7 and Fig. S8).

### Atlas is an essential transition protein

Several lines of evidence suggest that Atlas is a transition protein that facilitates the change from histone-based to protamine-based chromatin packaging in spermatid nuclei. Atlas localizes throughout spermatid nuclei (Fig. 6) and has biochemical properties consistent with direct DNA interaction. The protein appears specifically at the late canoe stage of nuclear compaction (Fig. 5B-D). Its lack of overlap with testis-specific histones (Fig. 5B), partial overlap with Mst35Bb (Fig. 5D), removal from needle-stage nuclei (Fig. 6) and absence from mature sperm (Figs. 5-6) are all consistent with the expression profile of a transition protein. Several other transition proteins have been characterized in *D. melanogaster*, including Tpl94D, thmg-1, thmg-2, and Mst84B [53, 58, 59]. Collectively, the transition proteins vary in the stage of nuclear condensation at which they first appear and the range of nuclear shapes over which they are found [79], but otherwise match Atlas in their biochemical properties, transient expression, and localization throughout the nucleus. Compared to these other transition proteins, Atlas is present over a fairly narrow range of nuclear condensation stages and reaches its peak expression just prior to the onset of individualization. *Atlas* is also the only transition protein gene characterized to date whose removal disrupts fertility, as *Tpl94D*, *thmg-1*, *thmg-2* and *Mst84B* mutants are all fertile [53, 58, 59]. This may reflect the relatively later timing of Atlas’s expression in spermatid nuclei, reduced functional redundancy between DNA-binding proteins at the later stages of condensation, a potential interaction between Atlas and an essential protamine-like protein, and/or a more stringent requirement for DNA binding at these stages.

Transition proteins give way in spermatid nuclei to protamine-like proteins, which bind DNA in mature sperm and persist through fertilization. In this way, protamine-like proteins function analogously to vertebrate protamines, though they are believed to be evolutionarily independent [57, 64]. In *D. melanogaster*, protamine-like proteins include Mst35Ba, Mst35Bb, Prtl99C and Mst77F [63, 80, 81]. Interestingly, while all characterized protamine-like proteins are present in mature sperm, only some are essential for fertility. Knockouts of *Mst35Ba*, *Mst35Bb*, or both show occasional nuclear shaping defects, but male fertility is normal [80, 82]. In contrast, mutants of *Prtl99C* or *Mst77F* are sterile. Prtl99C and Mst35Ba/b bind condensed DNA independently of each other and contribute additively to the shortening of needle-stage nuclei, but Prtl99C’s effect is ∼3-fold greater [63, 64]. This difference is apparently great enough to reduce fertility only in *Prtl99C* mutants. In contrast, *Mst77F* and *Mst35Ba/b* show a genetic interaction, as *Mst35Ba/b* null flies become nearly sterile in an *Mst77F* heterozygous background [81]. Furthermore, while Mst35Bb-GFP is expressed in *Mst77F* nulls, these flies show deformed spermatid nuclei that do not reach a recognizable needle-like stage. Because *atlas* nulls show considerable phenotypic similarity to *Mst77F* nulls, but not *Prtl99C* nulls, we hypothesize that Atlas may act in a pathway with Mst77F. Our observation of inefficient IC movement down sperm tails in *atlas* null testes is reminiscent of a similar phenotype in *Mst77F* nulls [81], providing further evidence that these proteins may act in a common pathway. In both cases, ICs can form at misshapen canoe-stage nuclei, but fail at a subsequent step. While the exact relationship between nuclear abnormalities in the late canoe stage and individualization is not entirely understood, it is possible that nuclear shape and the organization of nuclear bundles impact the ability of IC association and IC progression, as is also observed in mutants of another gene, *dPSMG1*, which controls nuclear shape [59].

### Evolution of atlas and other de novo genes

Because *de novo* genes emerge from non-coding sequences, they typically encode proteins that are short and lack complex structure [3, 16]. Indeed, expanding the length of the protein-coding region and evolving higher-level protein structures are hypothesized to be among the final stages of new gene evolution [4]. In light of these constraints, what kinds of cellular functions might be available to newly evolved proteins? Vakirlis et al. [83] overexpressed emerging proto-genes in *S. cerevisiae* and found that those encoding proteins with predicted transmembrane (TM) domains were more likely to be adaptive, as assessed by the effect of proto-gene overexpression on growth rate. Such proteins may arise when thymine-rich intergenic regions undergo mutations that allow protein-coding gene birth and expression, since many codons with multiple U nucleotides encode amino acids commonly found in TM domains [83]. Our imaging data, in addition to the prediction tools employed by Vakirlis et al. [83], suggest that Atlas does not contain a TM domain. However, just as the amino acid compositional requirements of a TM domain are not overly complex, neither are those of DNA binding proteins. In essence, these proteins must simply be small, have a high concentration of positively charged residues, and contain a nuclear localization signal, which itself requires a small patch of positively charged residues [65]. Thus, DNA binding proteins may be a relatively easy class of protein to evolve *de novo*.

While many putative *de novo* genes are expressed in the *D. melanogaster* testis [Fig. 1A and ref. 19], *atlas* was the only verified hit from our screen that was essential for male fertility. This result raises two related questions. First, why are so many of these other genes expressed if their knockdown causes no obvious effect? In general, it is common for the knockdown of protein-coding genes expressed in reproductive tissues in *D. melanogaster* to result in no detectable fertility defects [46, 84–86]. One set of explanations center on technical issues: a gene could be expressed outside of the cells targeted by the RNAi driver, knockdown level could be insufficient to cause a phenotype, and/or a gene may be functionally redundant such that its knockdown causes no apparent effect. Another hypothesis to explain this pattern is that while the loss of function of such genes may cause small reductions in fertility that would be subject to strong negative selection in nature, the conditions used to assay such knockdown animals in primary screens are rarely tailored to detect differences of this magnitude. A third possibility, not mutually exclusive with the above, is that while some genes may be expendable in non-competitive, non-exhaustive mating conditions, their absence may result in lower fitness in sperm competitive environment, environments in which males mate several times in quick succession, or environments in which sperm must persist in storage for longer intervals or during less optimal conditions [87–89]. These latter two possibilities are illustrated by some of the other spermatid chromatin binding proteins previously characterized as “non-essential” (e.g., *Mst35Ba/Bb* and *Tpl94D*) [58, 80]. While these proteins are not *de novo* evolved, they provide examples of genes whose mutations cause aberrant cellular phenotypes, such as the abnormally shaped spermatid nuclei in *Mst35Ba/Bb,* and that may contribute to fertility defects when placed in a sensitized genetic background [81].

It is also possible that some of the genes we screened are *de novo* genes that have no functional effects and have thus never been selectively advantageous. For example, 8 of the 42 screened genes have annotated orthologs in either *D. melanogaster* alone or in only *D. melanogaster* and *D. simulans*. Such recently “born” *de novo* genes could have been propagated by neutral evolutionary processes but not yet experienced inactivating mutations that would abolish their coding sequences and/or expression. This pattern would be consistent with *de novo* genes’ high rates of both birth and death observed previously in *Drosophila* [38] and yeast [83].

A second question raised by our finding that *atlas* encodes an essential transition protein is: how might *atlas* have evolved to become essential for fertility in *D. melanogaster*, particularly when other transition proteins appear functionally redundant? Other proteins involved in spermatid chromatin compaction show variable levels of conservation across *Drosophila*. For example, the protamines that contribute to DNA packaging in mature sperm [57] are found across all sequenced *Drosophila* species [61], and orthologs are also reported in FlyBase from other Dipteran and non-Dipteran insects [90]. However, transition protein Tpl94D is reported to be restricted to species ranging from *D. melanogaster* to *D. pseudoobscura* [60], as are the related proteins tHMG1 and tHMG2 with high-mobility group domains [58, 90]. Results like these suggest that while some protamine-like proteins (i.e., Mst35Ba and Mst35Bb) have consistently been among the final chromatin-packaging proteins, the specific proteins facilitating the transition from histones to protamines have likely varied over evolutionary time. Against this backdrop, and based on our analyses of the protein’s presence/absence, biochemical properties, and expression patterns in extant species, we hypothesize that while the Atlas protein likely had some DNA-binding ability and male-specific expression upon its origin, it was only one of several proteins involved in spermatid chromatin compaction. Since *atlas* was lost independently in several lineages after its birth (Fig. 7), Atlas was likely non-essential at its outset, but rather evolved an essential function within the *melanogaster* group of species. Such evolution of essentiality could have occurred because of the loss of a protein with a complementary function and/or changes in the process of spermiogenesis that thrust Atlas into a functionally unique role. It is also worth noting that species that have evidently lost *atlas* might have undergone other compensatory changes in their repertoires of spermatid DNA binding proteins. For instance, *D. willistoni* lacks *atlas* but appears to have several additional paralogs of the protamines found only in duplicate in *D. melanogaster*, which could have evolved transition-protein-like roles.

While our study cannot establish whether the movement of the *atlas* protein-coding sequence off of the X chromosome onto an autosome early in the evolution of the *melanogaster* group (Fig. 7) affected the gene’s essentiality, such movement remains noteworthy. Prior work has found a significant dearth of testis-expressed genes on the X chromosome in *Drosophila* [69, 91–93] and other species [94, 95]. Furthermore, *Drosophila* exhibit suppression of X-linked testis-expressed genes, and transfer of such genes from the X chromosome to autosomal loci results in higher expression levels [96–98]. One of several proposed mechanisms for both the paucity of X-linked testis-expressed genes and the suppression of their expression is meiotic sex chromosome inactivation (MSCI), in which the X chromosome becomes transcriptionally silenced earlier than autosomes [99–104]. Thus, genes that affect meiotic or post-meiotic processes, as *atlas* does, could exert beneficial effects more strongly and/or for a longer period of time if they become encoded autosomally. While the *atlas* protein-coding sequence appears to show male-specific expression regardless of its chromosomal location, it is possible that the movement of *atlas* to chromosome 2 allowed it to evolve a broader or different expression pattern that expanded or modified its role in spermiogenesis. The complex molecular bases of both X suppression and “escape” from the X chromosome in *Drosophila* continue to be actively investigated and debated [102–108], but continued research in this area might inform further interrogation of the forces driving *atlas* off of the X chromosome.

The movement of the *atlas* protein-coding sequence to chromosome 2 also created the two-exon gene observed in *D. melanogaster*, in which the longer second exon appears to be entirely non-coding. This second exon is highly conserved across the genus in both sequence and genomic location, and it shows male-specific expression in several species that lack the protein-coding sequence upstream (Fig. 7 and Fig. S8). These patterns of conservation suggest that the second exon might originally have been a non-coding RNA, a class of molecule whose importance in *Drosophila* male reproduction has recently become recognized [109, 110]. While previous examples of functional ncRNAs in spermatogenesis have generally acted in *trans* to regulate other genes or affect the functions of other proteins, it is also possible that the long 3’ UTR of *atlas* in *D. melanogaster* could affect the translation of *atlas* transcripts. Many genes functioning in spermatid differentiation are transcribed early in spermatogenesis but translationally repressed until later in spermiogenesis, a phenomenon that relies on various forms of post-transcriptional regulation [111, 112]. Future studies of the *atlas* protein-coding sequence in the absence of its 3’ UTR, the expression patterns of Atlas protein in species in which it is encoded from the X chromosome, or the genetic ablation of the conserved region in species lacking the protein-coding sequence will provide additional insights.

A final issue raised by our results is the exact timing and mechanism of origin for the *atlas* protein-coding sequence. The bioinformatic screen [19] that identified *atlas* and the other genes tested in Fig. 1 was designed to identify both “*de novo*” genes, defined as protein-coding regions in *Drosophila* that had recognizable, but non-ORF-maintaining, TBLASTN hits in outgroup species, and “putative *de novo*” genes, which had no TBLASTN hits in outgroup species. (Importantly, the screen also eliminated any protein with an identifiable protein domain, thus reducing the chances of identifying divergent members of gene families.) The vast majority of the genes we tested with RNAi, including *atlas*, fell into the putative *de novo* category. The bioinformatic screen’s criteria were reasonable for a high-throughput analysis, but BLAST-based methods have known limitations for detecting orthologous sequences in diverged species [74, 113, 114]. The lack of identifiable *atlas* protein-coding genes in several *Drosophila* species (e.g., *D. pseudoobscura* and *D. willistoni*) is unlikely to be due to BLAST homology detection failure, and extensive synteny-based searches confirmed the gene’s absence (Fig. S9). BLAST and synteny-based searches for orthologs in non-*Drosophila* species also did not detect an ortholog, though BLAST searches are not predicted to have adequate sensitivity for a protein of this size and evolutionary rate, at this level of species divergence [74]. Hence, in addition to using synteny to search for orthologs, we used HMMER, which employs hidden Markov models and builds a sequence profile of the target protein using information from multiple orthologs. Since HMMER also did not detect orthologs outside of *Drosophila*, we hypothesize that *atlas* evolved *de novo* at the base of the genus. However, since we remain unable to identify the non-protein-coding sequence from which *atlas* arose, we continue to refer to *atlas* as a putative *de novo* gene [5].

Overall, we find that while many putative *de novo* evolved genes are expressed in the *D. melanogaster* testes, few have major, non-redundant effects on fertility. However, several such genes have evolved critical roles at distinct stages of spermatogenesis and sperm function. We showed previously that the putative *de novo* gene *saturn* is required for maximal sperm production, as well as for the ability of transferred sperm to migrate successfully to sperm storage organs in females [46]. Another putative *de novo* gene, *goddard*, is required for sperm production and encodes a cytoplasmic protein that appears to localize to elongating axonemes [20, 46]. Loss of *goddard* impairs the individualization of spermatid bundles [20], thus exerting an effect that appears to be upstream of those observed for *saturn* and *atlas*. Here, we report another novel function for a putative *de novo* gene: encoding an essential transition protein that is necessary for proper nuclear condensation in spermiogenesis. Taken together, these results demonstrate that while many *de novo* genes may play subtle roles or share functional redundancy with other genes, *de novo* genes can also become essential players in complex cellular processes that mediate successful reproduction.

## Materials and Methods

### RNA Interference Screen

*De novo* and putative *de novo* genes inferred to be no older than the *Drosophila* genus were identified previously [19]. We filtered these genes with publicly available RNA-seq data [115] to identify those expressed predominantly in the testes [>50% of RPKM sum deriving from the testes from ModENCODE data; ref. 115], giving a total of 96 genes. To assess each of these candidates for effects on male fertility, we induced knockdown in the male germline by crossing UAS-RNAi flies to Bam-GAL4, UAS-Dicer2 flies [46, 116]. Control flies were generated by crossing the attP-containing genetic background into which UAS-RNAi was inserted to the same GAL4 line. Flies carrying UAS-RNAi were of two types. Roughly half of the genes had publicly available lines from the Vienna *Drosophila* Resource Center [117] or the Transgenic RNAi Project [50]. For the other genes, no publicly available RNAi stock was available, so we constructed TRiP-style stocks in the pValium20 vector as previously described [85]. These constructs were integrated into an AttP site in stock BL 25709 (*y*^1^ *v*^1^ P{nos-phiC31\int.NLS}X; P{CaryP}attP40) from the Bloomington *Drosophila* Stock Center (injections by Genetivision; Houston, TX, USA) and crossed into a *y v* background to screen for *v*^+^. We attempted at least two rounds of transgenic production for each gene. In total, we were able to obtain and test RNAi lines for 57 of the 96 identified genes. Table S3 shows all RNAi lines used and lists the short hairpin sequences cloned for the TRiP lines we constructed.

We initially screened males knocked down for each candidate gene for major fertility defects by crossing groups of 7 knockdown or control males to 5 virgin Canton S females, letting the adults lay eggs for ∼48 hours, and then discarding adults and quantifying the resulting progeny by counting the pupal cases, as previously described [46]. To assess the degree of knockdown achieved, 10 whole males of each line were homogenized in TRIzol reagent (Life Technologies, Carlsbad, CA). RNA isolation, DNAse treatment, cDNA synthesis and semi-quantitative RT-PCR with gene-specific primers were performed as previously described; amplification of *RpL32* was used as a positive control [46]. We evaluated knockdown efficiency by agarose gel electrophoresis of RT-PCR products as one of four levels: “complete” if no product from the knockdown cDNA sample was visible via agarose gel electrophoresis; “near complete” for a very faint knockdown product that was also much less abundant than the control product; “partial” for a more robust knockdown product that was still visibly less intense than control; and “not knocked down” if the product intensity for the knockdown sample equaled or exceeded that of the control. Any gene that did not show at least partial knockdown was discarded from further analysis, leaving a total of 42 genes successfully screened. Table S3 shows the degree of knockdown achieved for each line.

### CRISPR Genome Editing

To validate RNAi results for *atlas*, *CG43072* and *CG33284*, we used CRISPR/Cas9 genome-editing to generate null alleles that could be used for further analysis, as described previously [20]. Briefly, our general strategy was to design gRNAs in the pU6.3 vector (Drosophila Genome Resource Center (DGRC) #1362) that targeted each end of a locus. These plasmids, along with plasmids encoding gRNAs that targeted the *w*+ locus, were co-injected by Rainbow Transgenics (Camarillo, CA) into embryos laid by *vasa*-Cas9 females in a *w*+ background, Bloomington stock #51323 [118]. G_0_ animals were crossed to *w-* flies, and members of G_1_ broods with a higher-than-expected fraction of *w*-progeny were individually crossed to an appropriate balancer line and then PCR-screened for the desired deletion of the targeted locus.

We also constructed three frameshift, expected loss-of-function alleles for *atlas* by using CRISPR to induce non-homologous end joining at a single PAM site just downstream of the *atlas* start codon. *Vasa*-Cas9 embryos were co-injected and screened for w-progeny as described above. We then used squish preps to isolate DNA from G1 flies and used a PCR-RFLP assay to detect mutations. PCR products spanning the gRNA-targeted site were digested with *Bfa*I (New England Biolabs (NEB), Ipswich, MA); undigested products in which the expected *Bfa*I site was lost indicated a mutation, which was balanced and then confirmed by PCR and sequencing of homozygous mutant lines.

We used scarless CRISPR editing and homology-directed repair (HDR) to insert the GFP protein-coding sequence in-frame at the end of the *atlas* protein sequence (see Fig. S6; (https://flycrispr.org/scarless-gene-editing/) [66–68]. We first generated an *atlas*-GFP DNA construct by cloning the *atlas* protein-coding sequence into pENTR and using LR Clonase II (Thermo Fisher Scientific, Waltham, MA) to recombine the sequence with pTWG (DGRC #1076; T. Murphy), generating a C-terminally tagged *atlas*:GFP construct. We amplified the *atlas* fragment from *vasa*-Cas9 strain #51323 genomic DNA. Once *atlas*-GFP was obtained in a plasmid, we amplified it with primers that contained 5’ *Esp*3I sites and overhangs designed for Golden Gate Assembly (GGA) and that, in the case of the reverse primer, also added on 42 nucleotides downstream of the *atlas* stop codon to reach a PAM site identified by FlyCRISPR TargetFinder [119] as being optimal for Cas9/gRNA recognition and cleavage. The primer also introduced a mutation in the PAM site so that insertion of the designed piece of DNA into the genome *in vivo* would not be subject to re-cutting. We also used the NEB Q5 Site-Directed Mutagenesis kit to introduce a silent mutation into the *atlas* protein-coding sequence to eliminate an internal *Esp*3I site. The resulting construct was used as the “left” homology arm for homology-directed repair (HDR) editing. We constructed a “right” homology arm by using NEB Q5 PCR to amplify a 982-bp fragment downstream of the PAM site, using primers modified to contain *Esp*3I sites and overhangs compatible with GGA. We performed GGA by combining these left and right arms, a plasmid containing a PiggyBac transposase-excisable 3xP3-dsRed flanked by *Esp*3I sites, and backbone plasmid pXZ13, with *Esp*3I and T4 DNA ligase (NEB). A combination of colony PCR, restriction digestion and sequencing identified properly assembled plasmids suitable for HDR.

*Vasa*-Cas9 embryos were co-injected with the assembled plasmid and a pU6.3 plasmid encoding a gRNA targeting the region just downstream of the *atlas* stop codon. G0 flies were crossed to *w*^1118^ adults, and G1 flies were screened for red fluorescent eyes using the NIGHTSEA system (NIGHTSEA LLC, Lexington, MA). Six balanced lines from two independent G1 broods were established. To remove the dsRed from the *atlas* locus, we crossed these lines to a PiggyBac transposase line (BDSC #8285) and then selected against pBac and dsRed in the following generation. PCR and sequencing confirmed the expected “scarless” insertion of GFP at the *atlas* locus.

### atlas *Genomic Rescue Line*

We constructed an HA-tagged *atlas* rescue line that contained the *atlas* gene flanked by 1345 bp of sequence upstream of the start codon (but excluding the coding sequence of upstream gene *CG3124*) and 3000 bp of sequence downstream of the stop codon (including the full 3’ UTR) as follows. Genomic sequences were PCR amplified using Q5 High fidelity Polymerase (NEB), purified Canton S genomic DNA (Gentra Puregene Tissue Kit, Qiagen, Germantown, MD), and the atlas rescue F1/R1 and atlas rescue F3/R3 primer sets (see Table S4). The 3x-HA tag was likewise amplified from pTWH (DGRC 1100; T. Murphy) using atlas rescue F2/R2 primers. These DNA fragments were subsequently assembled into a XbaI/AscI-linearized w+attB plasmid (Addgene, Watertown, MA, plasmid 30326, deposited by J. Sekelsky). The assembled construct was then phiC31 integrated into the PBac{*y*^+^-attP-3B}VK00037 (Bloomington *Drosophila* Stock Center (BDSC) stock #24872) docking site (Rainbow Transgenics) and crossed into the *atlas* null background to assess rescue.

### Fertility Assays and Sperm Visualization

To validate the finding of reduced fertility for *atlas* knockdown males in the group fertility assay described above, we performed single-pair fertility assays in which knockdown or mutant males or their controls were mated individually to Canton S virgin females. Based on previous experience analyzing genes that resulted in sterility or near-sterility [20, 46], we designed assays with *N* = 20-30 flies per male genotype. Matings were observed, and males were discarded after copulation. Females were allowed to lay eggs into the vials for 4 days and then discarded. Pupal cases were counted as a measure of fertility. Crosses to generate and mating assays involving RNAi flies were maintained at 25° to optimize knockdown. Before all assays, flies were reared to sexual maturity (3-7 days) in single-sex groups on cornmeal-molasses food supplemented with dry yeast grains [46].

To assess the level of fertility conferred by the *atlas*-GFP allele, we crossed *atlas*-GFP and *w*^1118^ flies to Δ*atlas*/SM6. Males with genotypes atlas-GFP/Δ*atlas* and +/Δ*atlas* were compared using the single-pair fertility assay described above.

To observe the production of sperm in knockdown or mutant males, we introduced the Mst35Bb-GFP marker into these males, which labels mature sperm and late-stage spermatid nuclei with GFP [51]. Samples were prepared, imaged and analyzed as described previously [46].

### Atlas-GFP Ectopic Expression

We used the Gateway cloning system (Thermo) to construct an *atlas*-GFP transgene expressed under UAS control (primers in Table S4). The *atlas* protein-coding sequence in pENTR was recombined with pTWG (*Drosophila* Genomics Resource Center, T. Murphy) as described above. The resulting plasmid was then inserted into w-flies using P-element-mediated transposition (Rainbow Transgenics), w+ G1s were selected, and several independent insertions were balanced. We crossed male UAS-*atlas*:GFP flies to females from two different driver lines: *tubulin*-GAL4 (to drive ubiquitous expression) and *Bam*-GAL4 (to drive expression in the early germline). We dissected larval salivary glands of the *tub*>*atlas*:GFP males, since these cells are exceptionally large and ideal for visualizing subcellular localization. We then dissected the testes of *Bam*>*atlas*:GFP males to evaluate whether the localization pattern observed in the salivary gland was consistent in testis tissue, albeit not the same cells in which endogenous atlas appears to be expressed. Protein localization was visualized by fluorescence confocal microscopy on a Leica SP5 microscope (Leica Microsystems, Wetzlar, Germany) and images were captured with LASAF as described previously [20].

### Imaging Spermatogenesis and Spermatid Nuclear Condensation

We used phase-contrast microscopy to examine the stages of spermatogenesis in whole mount testes [120]. To assess the processes of nuclear condensation and individualization of 64-cell cysts of spermatids in the post-meiotic stages of spermatogenesis, we used fluorescence and confocal microscopy to visualize actin-based individualization complexes and nuclei. Samples were processed, and actin and nuclear DNA were visualized with TRITC-phalloidin (Molecular Probes, Eugene, OR) and TOPRO-3 iodine (Thermo), respectively, as described previously [20]. The final stages of nuclear condensation were visualized with the Mst35Bb-GFP marker described above, as well as an equivalent marker, Mst35Bb-dsRed [51]. Earlier nuclear stages were visualized with histone H2AvD-RFP (BDSC stock #23651), which is present in round spermatid nuclei and the earliest stages of nuclear elongation [53, 121].

Images with H2AvD-RFP were obtained with epifluorescence microscopy, since we lack an appropriate confocal laser for RFP. To examine spermatid nuclei at various stages of condensation, we visualized nuclear bundles using TOPRO-3. Testes of newly eclosed (<1 day old) *atlas* null and control males were dissected in PBS. Testes were then transferred to a droplet of 2% paraformaldehyde on poly-L-lysine treated glass slides and were gently shredded in the post-meiotic region to release sperm bundles. Testes were gently squashed beneath coverslips coated in Sigmacote (Sigma Aldrich, St. Louis, MO). We then froze slides in liquid nitrogen for a few seconds and popped off of the siliconized coverslip with a razor. Slides were incubated in Coplin jars filled with 95% ethanol at −20°C for 30 minutes and then mounted in VECTASHIELD (Vector Laboratories, Burlingame, CA). Nuclear staging was performed by examining the shape of the nuclei. Early and late canoe stages of condensation were distinguished by the absence or presence of Mst35Bb-GFP, respectively. Elongated and late canoe stages were distinguished by the presence or absence, respectively, of vesicles of GFP-tagged nuclear proteins (Atlas-GFP or Mst35Bb-GFP) located basal to the nuclei. Examples of stages are given in Fig. 4. Confocal stacks were taken on a Leica SP5 microscope, images were captured by LASAF, and ImageJ was used to flatten stacks into a single, two-dimensional image. All intact nuclear bundles were counted for each dissection.

For the experiments measuring nuclear condensation stage (Table S1), a sample size of *N* = 10 for each genotype was selected based on the magnitude of the *atlas* null phenotype and the consistent differences observed in previous dissections of these genotypes with Mst35Bb-GFP. Likewise, for the IC-nuclear bundle association and IC progression analysis (Fig. 3C-D), we selected sample sizes of *N* = ∼15 per genotype based on pilot experiments showing that aberrant actin phenotypes were highly consistent in null testes and previous experience with such quantification [20].

### Evolutionary and Gene Expression Analysis of atlas

We searched for orthologs of the *D. melanogaster* Atlas protein in the original 12 sequenced *Drosophila* species with BLASTP searches in FlyBase [122]. We also used TBLASTN searches to identify orthologs in species lacking complete protein annotations. We identified syntenic regions for each species by looking for conserved neighboring genes, such as *ord* and *CG3124*. In addition to analyzing the *atlas* coding region, we conducted separate BLASTN searches for the sequence of the *D. melanogaster* 3’UTR across *Drosophila* species since it has a different conservation pattern than the coding sequence.

To test for sex-specific expression biases for both the ORF and the 3’ UTR sequences, we used adult male- and female-specific RNA-seq data from numerous *Drosophila* species accessed through the Genomics Education Partnership version of the UCSC Genome Browser (http://gander.wustl.edu/) and initially collected by Brown et al. [115] and Chen et al. [123]. We also confirmed these findings experimentally in several species by performing RT-PCR on cDNA isolated from whole males and whole females, as previously described [46].

To search for *atlas* orthologs in non-*Drosophila* Dipterans, we obtained from ENSEMBL Metazoa the genomes of *Musca domestica, Glossina morsitans, Lucilia cuprina, Aedes aegypti, Anopheles darlingi, Anopheles gambiae, Culex quinquefasciatus* and *Mayetiola destructor*. We performed a synteny search (summarized in Fig. S11) in each species by identifying the nearest neighbors of *atlas* in the *D. ananassae* and *D. virilis* genomes that had an identifiable homolog in each species. In all cases, the homologs of the nearest neighbors on each side of *atlas* were found on different contigs, suggesting synteny breakdown. We obtained up to 1 Mb of sequence on each side of each identified homolog and queried it with BLASTN, TBLASTN, and Exonerate [124] for regions with significantly similarity to any portion of the Atlas protein or cDNA sequences. No significant hits, and no hits better than what could be found in other parts of the genome, were found. Finally, we used HMMER to search for orthologs in all annotated proteomes and all metazoan proteomes. We first queried the database with Atlas from either *D. melanogaster* or *D. virilis* and accepted hits that fell below an e-value cutoff of 0.01 and a minimum hit length of 3%. These hits were then included iteratively in subsequent searches until no new significant hits were found.

We analyzed the molecular evolution of the *atlas* protein-coding sequence by obtaining orthologous protein-coding sequences from *melanogaster* group species. (Analysis out of this group was not performed due to high sequence divergence and poor alignment quality.) We used BLASTP to identify these sequences from GenBank and then extracted the coding DNA sequence for each. Sequences were aligned, checked for recombination, used to construct a gene tree, and analyzed with the PAML sites test as described previously [78], except that alignment positions that included gaps were masked from the PAML analysis. We initially analyzed a set of 13 species (*melanogaster*, *simulans*, *sechellia*, *yakuba*, *erecta*, *suzukii*, *takahashii*, *biarmipes*, *rhopaloa*, *ficusphila*, *elegans*, *eugracilis* and *ananassae*); we excluded an ortholog detected in *D. bipectinata* due to poor alignment. This initial analysis detected a class of sites with significant evidence of positive selection, but closer inspection of the alignment revealed that the site with the strongest evidence of selection, corresponding to *D. melanogaster* residue 31R, may have been driven by a questionable alignment due to an insertion in that region that was unique to *D. takahashii*. The reported results come from an analysis that excluded *D. takahashii*, which produced a more reliable alignment and showed no evidence for any sites under positive selection.

## Acknowledgements

We thank Dr. Alexis Hill for assistance with developing the *atlas*-GFP transgene; Gynesis Vance, Elvis Perez, Ishanpepe Jagusah, Emily Gualdino, and students in the Fall 2016 and Fall 2017 sections of Biology 261 Lab at Holy Cross for assistance with RNAi fertility screens; Dr. Rob Bellin for assistance with confocal microscopy; Dr. Justin McAlister for use of the NIGHTSEA system; the Bloomington Stock Center and the Vienna *Drosophila* Resource Center for fly strains; and, Dr. Mariana Wolfner and members of the Findlay and Bornberg-Bauer labs for productive discussions about the project and the manuscript. This work was supported by NSF CAREER Award #1652013 (to GDF) and a Humboldt Fellowship (to AG). GDF and EBB are also grateful to the Evolution Think Tank at the University of Münster, which funded GDF as a visiting fellow for in-person collaboration.

