## Supplemental material for "A putative *de novo* evolved gene required for spermatid chromatin condensation in *Drosophila melanogaster*"

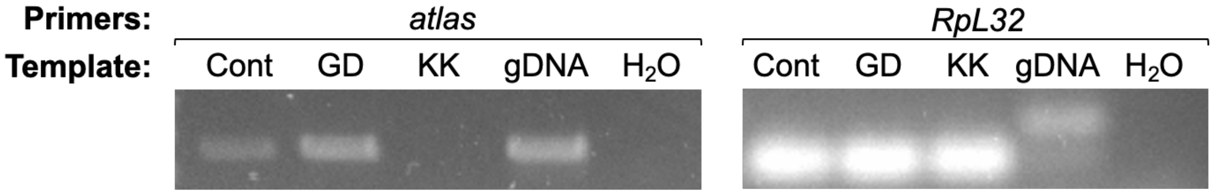

**Fig. S1: Example RT-PCR demonstrating near-complete knockdown of *atlas*.** Knockdown was driven by crossing strain VDRC KK-108680 to *Bam*-GAL4, UAS-*Dicer2*. We also attempted to induce knockdown in the same manner with strain VDRC GD-17240, and we produced control flies by crossing VDRC attP strain #60100 to *Bam*-GAL4, UAS-Dicer2. cDNA was isolated from whole males of each strain, and a standardized amount of cDNA or control genomic DNA from $w^{1118}$ was assessed for amplification of *atlas* and a housekeeping control gene, *RpL32*. The GD line did not induce detectable knockdown, but the KK line showed near-complete knockdown of *atlas*. Knockdown was assessed in the same way for all other RNAi lines tested; the control cross for TRIP-style RNAi lines was *y v 1509* crossed to *Bam*-GAL4, UAS-*Dicer2.*

A

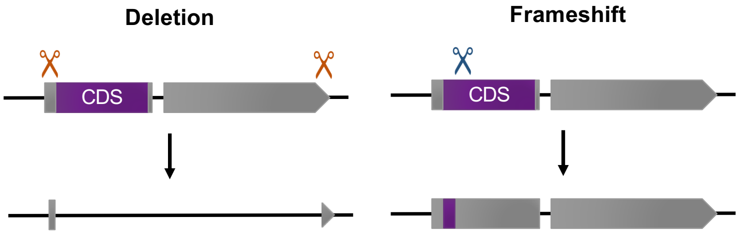

B

```
WT     GATAATAATATTATCAAAAACGTCAGAGTATTTATCGTATTGTATATTTCCAAATTACTT   60
atl52  GATAATAATATTATCAAAAACGTCAGAGTATTTATCGTATTGTATATTTCCAAATTACTT   60
atl62  GATAATAATATTATCAAAAACGTCAGAGTATTTATCGTATTGTATATTTCCAAATTACTT   60
atl86  GATAATAATATTATCAAAAACGTCAGAGTATTTATCGTATTGTATATTTCCAAATTACTT   60

WT     GTTATCACATCGAGTAAATGGGACGCAAAGGCCACAAGACTAGAA---------------  105
atl52  GTTATCACATCGAGTAAATGGGACGCAAAGGCCACAAGA-----AAAGGAAAGGCCTTCC  115
atl62  GTTATCACATCGAGTAAATGGGACGCAAAGGCCTTT------------------------   96
atl86  GTTATCACATCGAGTAAATGGGACGCAAAGGCCACAAGA-----A---------------  100

WT     ----------GGAAAGTTAAAGTCCACTCGAAGGTCCATCCTCCTCAGAAAATTAAAGCG  155
atl52  TTTTTAAGAAGGAAAGTTAAAGTCCACTCGAAGGTCCATCCTCCTCAGAAAATTAAAGCG  175
atl62  -----------GAAAGTTAAAGTCCACTCGAAGGTCCATCCTCCTCAGAAAATTAAAGCG  145
atl86  ----------GGAAAGTTAAAGTCCATTCGAAGGTCCATCCTCCTCAGAAAATTAAAGCG  150

WT     AAGCTGCCGAAAAGAGATAAACGTAGCACACTTCGTCCCAAGATTAATAATTCGAAGGTT  215
atl52  AAGCTGCCGAAAAGAGATAAACGTAGCACACTTCGTCCCAAGATTAATAATTCGAAGGTT  235
atl62  AAGCTGCCGAAAAGAGATAAACGTAGCACACTTCGTCCCAAGATTAATAATTCGAAGGTT  205
atl86  AAGCTGCCGAAAAGAGATAAACGTAGCACACTTCGTCCCAAGATTAATAATTCGAAGGTT  210

WT     TTTCACGTGAACTGCTGCAAGTACAAGATCCGTTCGAGACCCATGAGGCGCAATCAGTGC  275
atl52  TTTCACGTGAACTGCTGCAAGTACAAGATCCGTTCGAGACCCATGAGGCGCAATCAGTGC  295
atl62  TTTCACGTGAACTGCTGCAAGTACAAGATCCGTTCGAGACCCATGAGGCGCAATCAGTGC  265
atl86  TTTCACGTGAACTGCTGCAAGTACAAGATCCGTTCGAGACCCATGAGGCGCAATCAGTGC  270

WT     AACCTGGCCACCCAGTGCCCCTCCCACACGGTCGCCATCGCCCCGTCTGTGCCGCTCCTG  335
atl52  AACCTGGCCACCCAGTGCCCCTCCCACACGGTCGCCATCGCCCCGTCTGTGCCGCTCCTG  355
atl62  AACCTGGCCACCCAGTGCCCCTCCCACACGGTCGCCATCGCCCCGTCTGTGCCGCTCCTG  325
atl86  AACCTGGCCACCCAGTGCCCCTCCCACACGGTCGCCATCGCCCCGTCTGTGCCGCTCCTG  330

WT     CATCATCCTGTCCGGCGGAAGGTGGTCCATATGCCGGAAACCGTGGTCATGCACATGCGG  395
atl52  CATCATCCTGTCCGGCGGAAGGTGGTCCATATGCCGGAAACCGTGGTCATGCACATGCGG  415
atl62  CATCATCCTGTCCGGCGGAAGGTGGTCCATATGCCGGAAACCGTGGTCATGCACATGCGG  385
atl86  CATCATCCTGTCCGGCGGAAGGTGGTCCATATGCCGGAAACCGTGGTCATGCACATGCGG  390

WT     AATATGGAGGACGCCCGCCGAGTGGCGGTTAAGCACAGAAGAACCCTTATTCAAACCCAT  455
atl52  AATATGGAGGACGCCCGCCGAGTGGCGGTTAAGCACAGAAGAACCCTTATTCAAACCCAT  475
atl62  AATATGGAGGACGCCCGCCGAGTGGCGGTTAAGCACAGAAGAACCCTTATTCAAACCCAT  445
atl86  AATATGGAGGACGCCCGCCGAGTGGCGGTTAAGCACAGAAGAACCCTTATTCAAACCCAT  450

WT     GTTCCTTCGCCGGATGTGGATGATGACGAAGATACACACCAAATCGATCGGAGGTCAAGG  515
atl52  GTTCCTTCGCCGGATGTGGATGATGACGAAGATACACACCAAATCGATCGGAGGTCAAGG  535
atl62  GTTCCTTCGCCGGATGTGGATGATGACGAAGATACACACCAAATCGATCGGAGGTCAAGG  505
atl86  GTTCCTTCGCCGGATGTGGATGATGACGAAGATACACACCAAATCGATCGGAGGTCAAGG  510

WT     CAAGCCTATAGGATGGGTCAAATGAGACCCTTATCCAGCGACGTGGATGTGGATGACATG  575
atl52  CAAGCCTATAGGATGGGTCAAATGAGACCCTTATCCAGCGACGTGGATGTGGATGACATG  595
atl62  CAAGCCTATAGGATGGGTCAAATGAGACCCTTATCCAGCGACGTGGATGTGGATGACATG  565
atl86  CAAGCCTATAGGATGGGTCAAATGAGACCCTTATCCAGCGACGTGGATGTGGATGACATG  570

WT     GATGTGGACCCGTTTCAGTAG                                         596
atl52  GATGTGGACCCGTTTCAGTAG                                         616
atl62  GATGTGGACCCGTTTCAGTAG                                         586
atl86  GATGTGGACCCGTTTCAGTAG                                         591
```

C

```
WT     MGRKGHKTRRKVKVHSKVHPPQKIKAKLPKRDKRSTLRPKINNSKVFHVNCCKYKIRSRP   60
atl52  MGRKGHKKRKGLPF*---------------------------------------------   15
atl62  MGRKGL*-----------------------------------------------------    7
atl86  MGRKGHKKES*-------------------------------------------------   11

WT     MRRNQCNLATQCPSHTVAIAPSVPLLHHPVRRKVVHMPETVVMHMRNMEDARRVAVKHRR  120
atl52  ------------------------------------------------------------   15
atl62  ------------------------------------------------------------    7
atl86  ------------------------------------------------------------   11

WT     TLIQTHVPSPDVDDDEDTHQIDRRSRQAYRMGQMRPLSSDVDVDDMDVDPFQ          172
atl52  ----------------------------------------------------           15
atl62  ----------------------------------------------------            7
atl86  ----------------------------------------------------           11
```

**Fig. S2: CRISPR editing methods for constructing *atlas* loss-of-function mutations.** A) Graphical representation of CRISPR loss-of-function strategies. Purple boxes represent the *atlas* coding region, while gray represents non-coding regions of the gene. Scissors indicate locations where gRNAs targeted Cas9-mediated double-stranded breaks. The deletion allele was generated using two sgRNAs targeting either side of *atlas* to excise the complete coding region (CDS) and nearly all of the noncoding region. Frameshift alleles were created using one sgRNA targeting a cut at the start of the *atlas* coding region that was repaired with non-homologous end joining (NHEJ). B) Alignment of mutations generated in frameshift alleles. The gene's start codon is indicated with dark green shading. Blue shading indicates bases inserted by NHEJ, orange shading indicates NHEJ deletions. All three mutations consist of net insertions or deletions that are non-multiples of three, resulting in premature stop codons indicated with underlining. The mutant alleles retain the possibility of encoding a truncated form of Atlas protein if a downstream start codon (light green shading) is used, since this codon is in-frame with the sequence encoding the protein's C terminus. C) Predicted protein sequences encoded by the wild-type and frameshift alleles. Blue shading indicates novel amino acids created by NHEJ indel mutations. Gray shading indicates the potential truncated Atlas protein that could be encoded by the frameshift alleles if translation initiated at the downstream start codon indicated in panel B. Such hypothetical, N-terminally truncated forms of Atlas protein would contain amino acids 61-172 of the wild-type protein. Yellow shading indicates the putative nuclear localization signal.

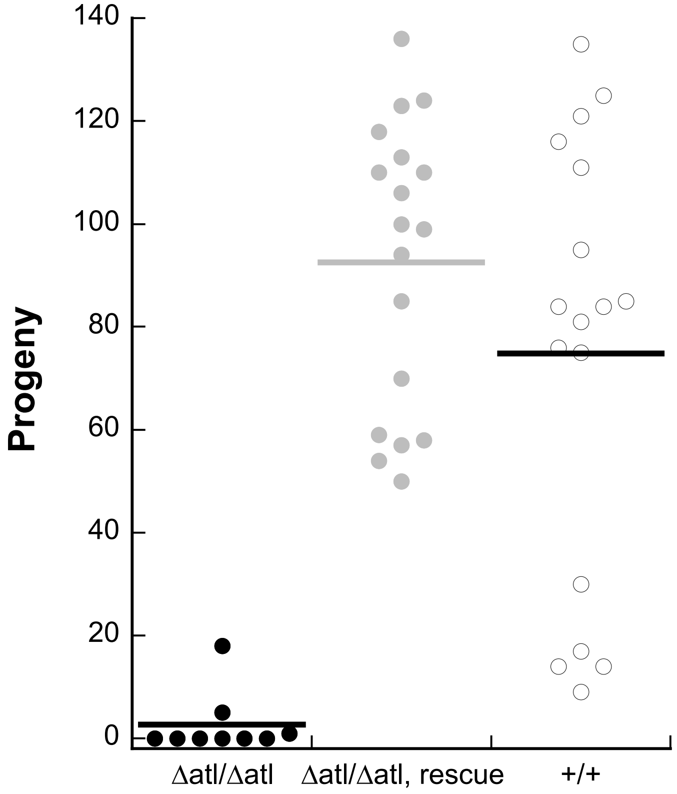

**Fig. S3: Atlas-HA genomic rescue construct restores full fertility in the *atlas* null background.** A single-pair fertility assay was used to assess fertility in males carrying a single copy of the *atlas*-HA genomic rescue construct in the *atlas* null background. These males showed no significant difference in fertility from control ($w^{1118}$) males (rescue fertility: 92.6 ± 6.6; control fertility: 74.8 ± 10.3; two-sample t-test assuming unequal variances, $p = 0.16$). Both control and rescue males had significantly higher fertility than *atlas* null males (null fertility: 2.7 ± 2.0; two-sample t-tests, both $p < 10^{-5}$).

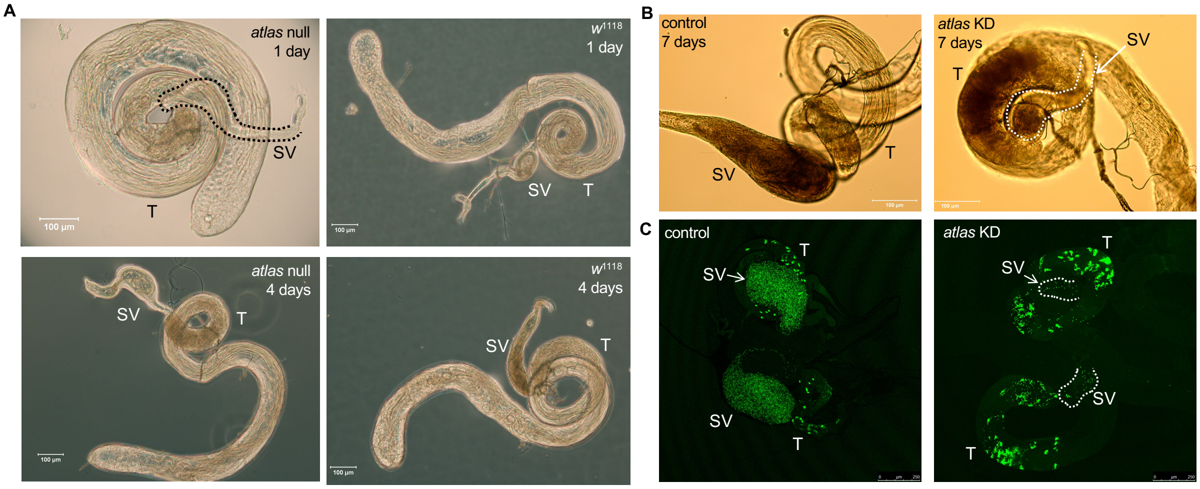

**Fig. S4: Accumulation of sperm in *atlas* null and knockdown testes.** A) Phase contrast imaging of *atlas* null and control males ages 1 day or 4 days. Sperm accumulate in the SV by 4 days in controls, but accumulate in the basal testes of null males on days 1 and 4. The day 7 images are shown in Fig. 3 of the main text. B) The same phenotype of sperm accumulation in the basal testis is observed in 7-day-old knockdown males. C) Knockdown males expressing Mst35Bb-GFP show similar patterns of spermatid nuclei to nulls, while control males accumulate many sperm nuclei in the SV (compare to Fig. 3 in the main text). Control flies in B-C were generated by crossing VDRC strain #60100 (attP) to Mst35Bb-GFP; + ; *Bam*-GAL4, UAS-*Dicer2*. SVs are highlighted for clarity when needed with dotted lines.

**Table S1: Stages of nuclear condensation observed in spermiogenesis in wild-type and *atlas* null testes.** Each line shows the distribution of staged nuclear bundles dissected from one individual pair of testes. Examples of nuclear stages and the curled nuclear phenotype observed in *atlas* null males are shown in Fig. 4.

| Sample | Normal round nuclear bundles | Normal early nuclear bundles | Normal late canoe nuclear bundles | Normal elongated nuclear bundles | Curled elongated nuclear bundles | Normal condensed nuclear bundles | Curled condensed nuclear bundles | % Curled nuclear bundles |
|---|---|---|---|---|---|---|---|---|
| WT 1 | 0 | 9 | 10 | 27 | 0 | 8 | 0 | 0% |
| WT 2 | 0 | 6 | 11 | 13 | 0 | 9 | 0 | 0% |
| WT 3 | 0 | 6 | 7 | 18 | 0 | 11 | 0 | 0% |
| WT 4 | 0 | 7 | 11 | 20 | 0 | 9 | 0 | 0% |
| WT 5 | 0 | 7 | 8 | 23 | 0 | 11 | 0 | 0% |
| WT 6 | 0 | 7 | 7 | 10 | 0 | 2 | 0 | 0% |
| WT 7 | 0 | 8 | 15 | 22 | 0 | 6 | 0 | 0% |
| WT 8 | 0 | 8 | 9 | 6 | 0 | 0 | 0 | 0% |
| WT 9 | 0 | 11 | 11 | 27 | 0 | 12 | 0 | 0% |
| WT 10 | 1 | 11 | 9 | 19 | 0 | 11 | 0 | 0% |
| Null 1 | 0 | 12 | 13 | 0 | 17 | 0 | 8 | 50% |
| Null 2 | 0 | 6 | 16 | 0 | 8 | 0 | 18 | 54% |
| Null 3 | 0 | 14 | 12 | 0 | 6 | 0 | 3 | 26% |
| Null 4 | 0 | 12 | 12 | 0 | 12 | 0 | 13 | 51% |
| Null 5 | 0 | 6 | 23 | 0 | 9 | 0 | 24 | 53% |
| Null 6 | 0 | 2 | 8 | 0 | 5 | 0 | 27 | 76% |
| Null 7 | 0 | 0 | 0 | 0 | 3 | 0 | 23 | 100% |
| Null 8 | 0 | 5 | 7 | 0 | 9 | 0 | 19 | 70% |
| Null 9 | 0 | 5 | 7 | 0 | 10 | 0 | 17 | 69% |
| Null 10 | 0 | 6 | 9 | 0 | 10 | 0 | 13 | 60% |

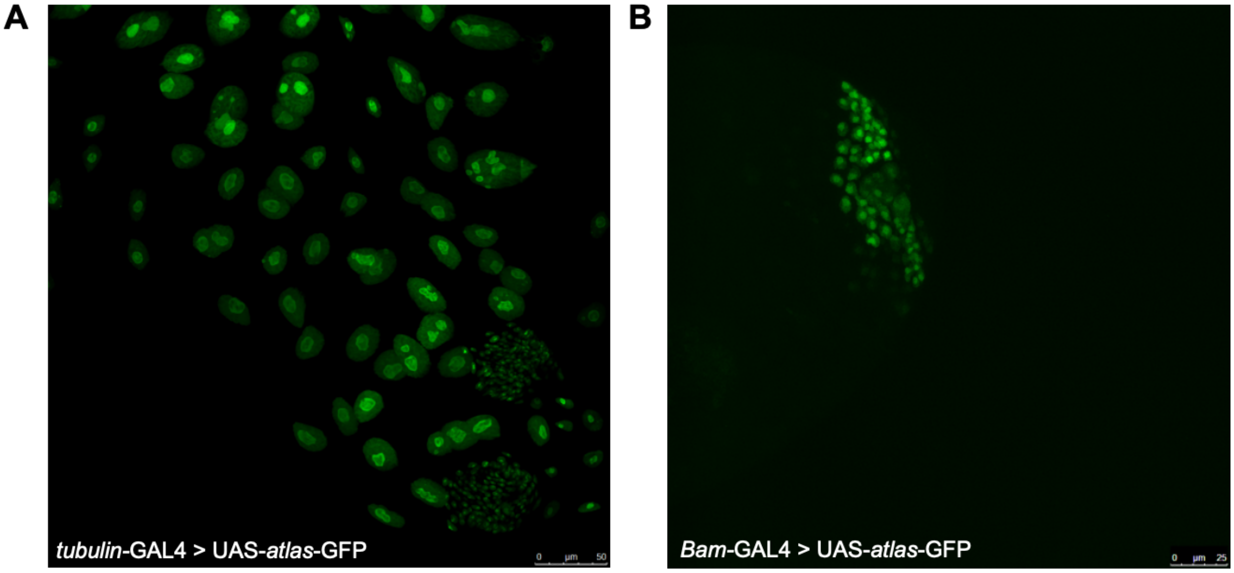

**Fig. S5: Expression of s*atlas*-GFP driven in larval salivary glands and early male germline cells.** A) Dissected larval salivary glands expressing UAS-*atlas*-GFP under the control of *tubulin*-GAL4. B) Apical portion of a testis expressing UAS-*atlas*-GFP under the control of *Bam*-GAL4. In both cases, Atlas-GFP has a predominantly nuclear localization pattern.

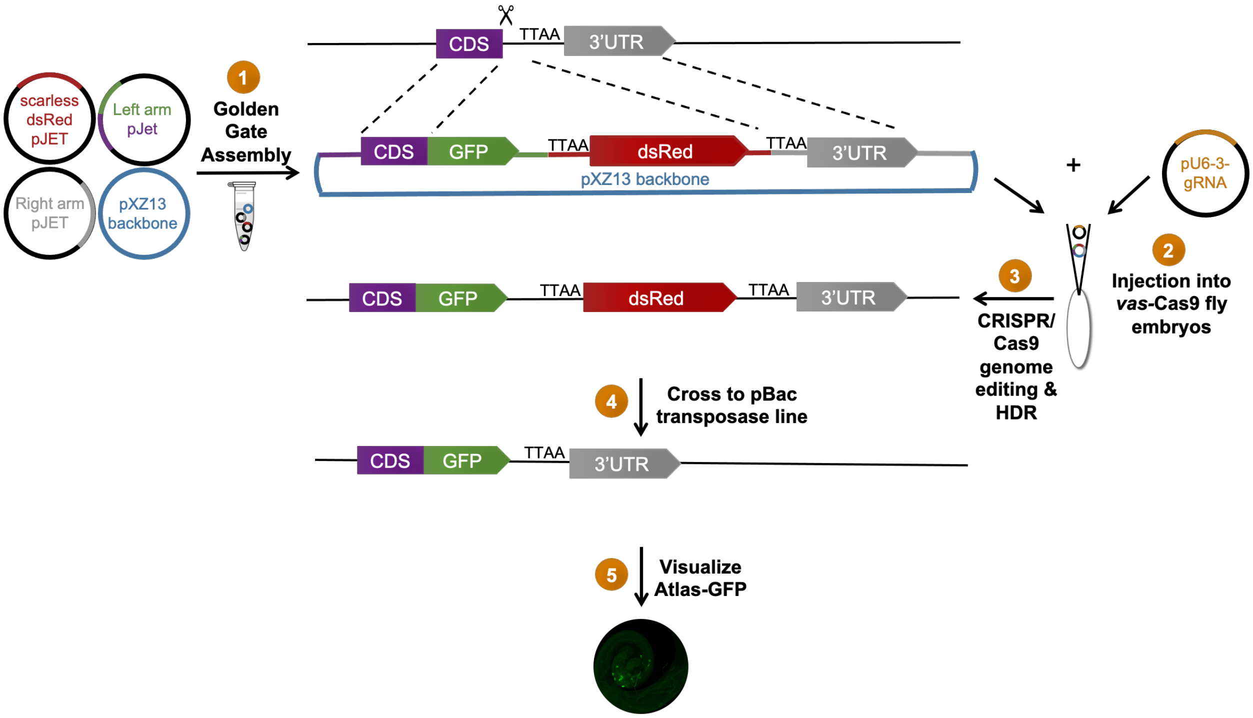

**Fig. S6: Scarless CRISPR/Cas9 genome editing strategy to produce a GFP knock-in allele at the endogenous *atlas* locus.** Golden Gate assembly was used to construct a plasmid carrying left and right homology arms flanking GFP placed in frame with the end of the *atlas* protein-coding sequence and a dsRed marker under the control of the 3xP3 promoter, which drives expression in the eye. This plasmid was injected into *vasa*-Cas9 flies along with a pU6.3 plasmid containing a gRNA targeting the end of the *atlas* coding sequence. $G_0$ flies were crossed to $w^{1118}$, and dsRed positive flies were screened molecularly for the correct *atlas*-GFP insert at the endogenous locus. The dsRed construct was then excised by crossing to a pBac transposase line, which removed the dsRed using flanking TTAA sequences. This protocol was adapted from Hill et al. (2019) and described at https://flycrispr.org/.

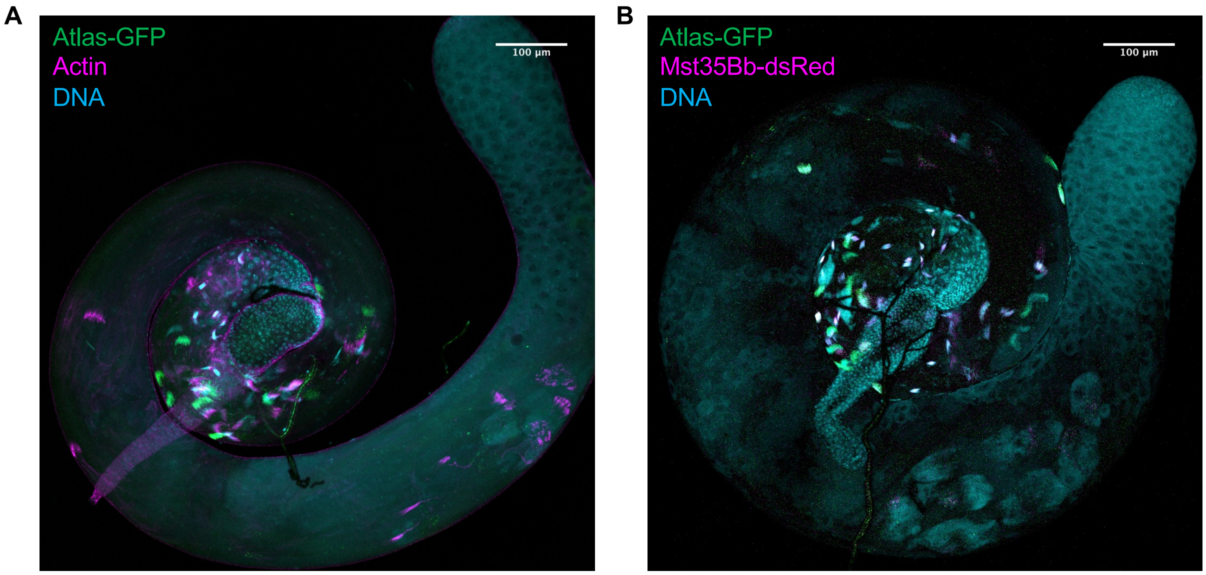

**Fig. S7: Images of whole testes from which the zoomed in basal portions in Figure 5 are taken.** A) Whole testis dissection corresponding to Figure 5C. Small GFP-positive puncta are visible near the progressed actin cones, which may represent the removal of Atlas-GFP from condensed nuclei. Also see Fig. 6D. B) Whole testis dissection corresponding to Figure 5D.

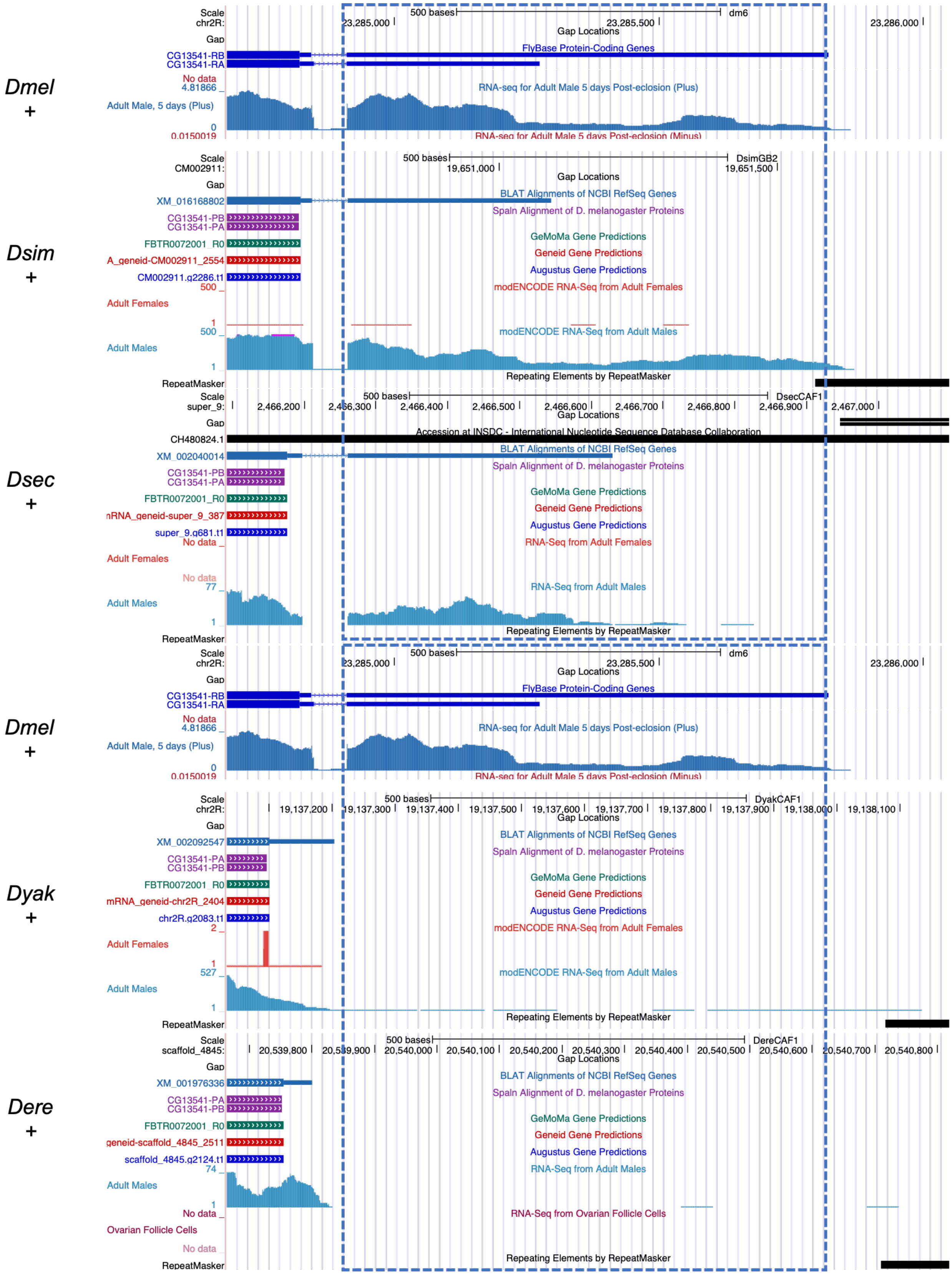
Dmel
+
Dsim
+
Dsec
+
Dmel
+
Dyak
+
Dere
+
FlyBase Protein-Coding Genes
RNA-seq for Adult Male 5 days Post-eclosion (Plus)
BLAT Alignments of NCBI RefSeq Genes
Spaln Alignment of D. melanogaster Proteins
GeMoMa Gene Predictions
Geneid Gene Predictions
Augustus Gene Predictions
modENCODE RNA-Seq from Adult Females
modENCODE RNA-Seq from Adult Males
Repeating Elements by RepeatMasker
RNA-Seq from Adult Females
RNA-Seq from Adult Males
RNA-Seq from Ovarian Follicle Cells

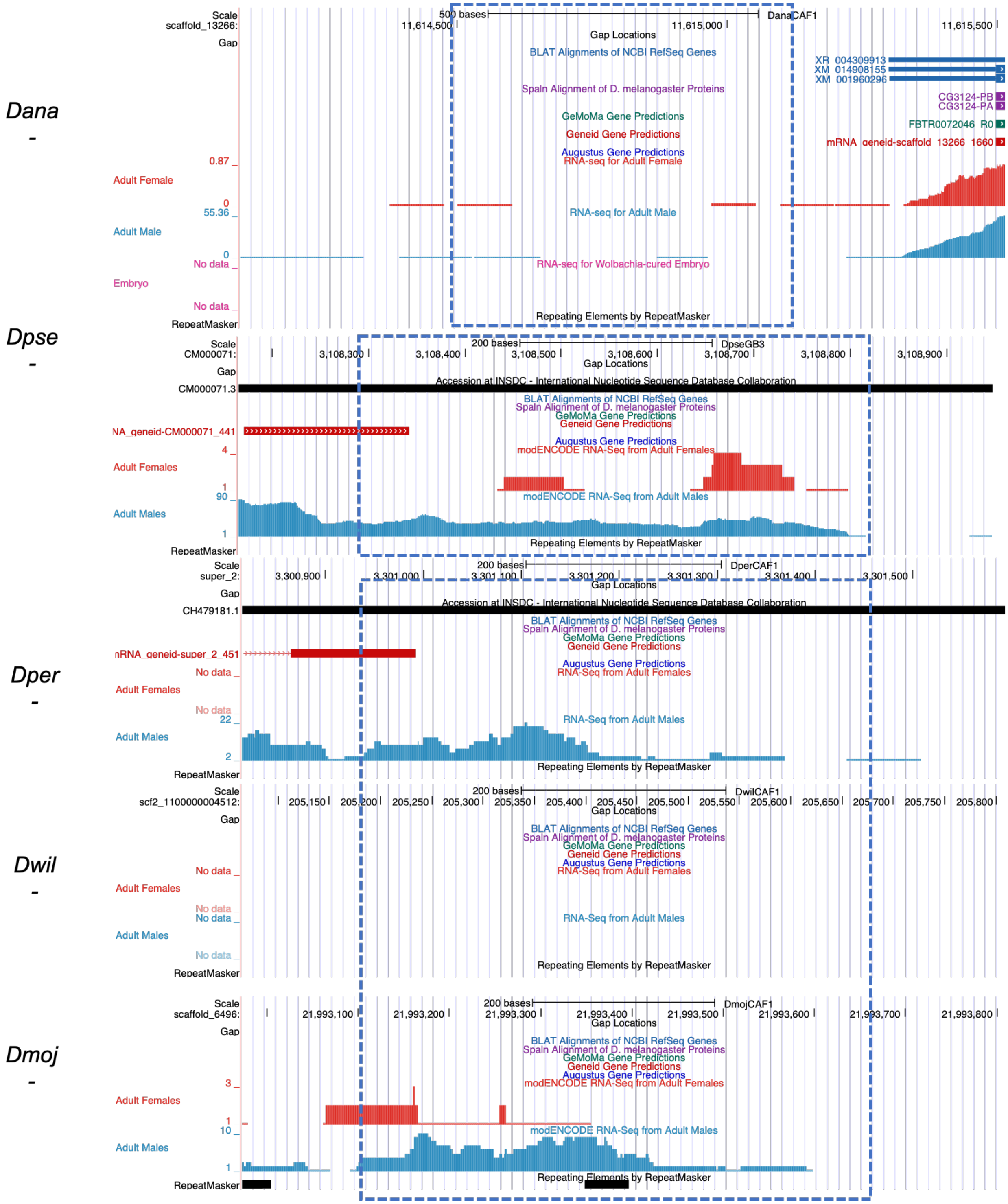
Dana
Dpse
Dper
Dwil
Dmoj
DanaCAF1
DpseGB3
DperCAF1
DwilCAF1
DmojCAF1
Gap Locations
BLAT Alignments of NCBI RefSeq Genes
Spaln Alignment of D. melanogaster Proteins
GeMoMa Gene Predictions
Geneid Gene Predictions
Augustus Gene Predictions
RNA-seq for Adult Female
RNA-seq for Adult Male
RNA-seq for Wolbachia-cured Embryo
modENCODE RNA-Seq from Adult Females
modENCODE RNA-Seq from Adult Males
RNA-Seq from Adult Females
RNA-Seq from Adult Males
Repeating Elements by RepeatMasker
Accession at INSDC - International Nucleotide Sequence Database Collaboration
Adult Female
Adult Male
Embryo
Adult Females
Adult Males
No data

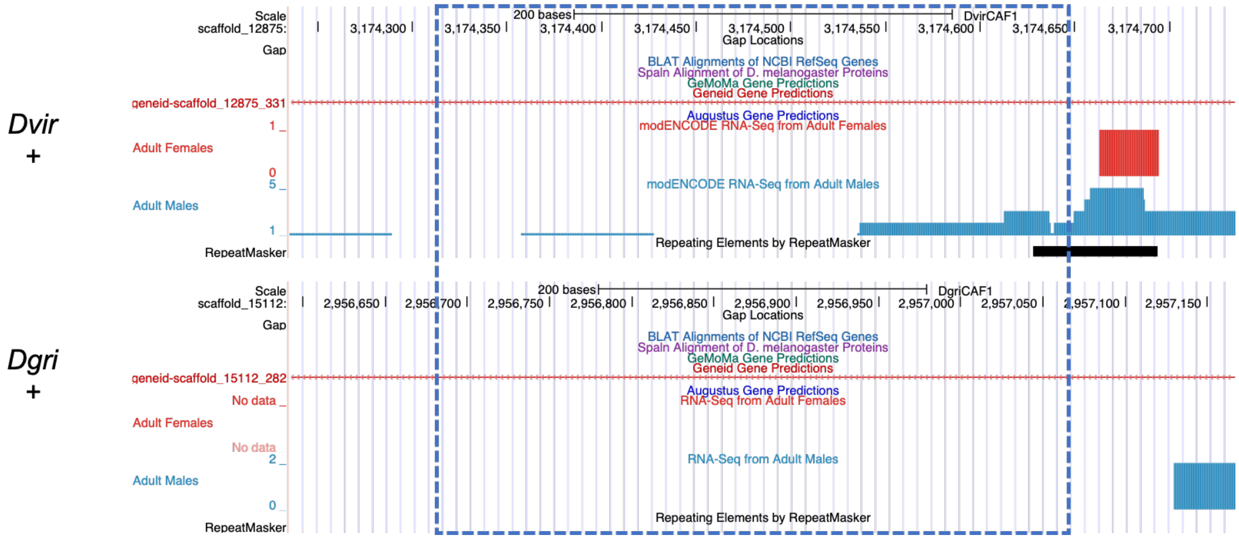

**Fig S8. The region of the genome corresponding to the non-coding exon of *atlas* in *D. melanogaster* is detectable across *Drosophila*, but expressed in only some species.** The non-coding second exon of *atlas* from *D. melanogaster* was compared with BLASTN to 11 other *Drosophila* species. The blue dashed line indicates the region that showed significant sequence identity in these searches. The Adult Male RNA-Seq track shows evidence of male-expressed RNA in the region. Peak heights are not comparable across species because the RNA-Seq was performed at different times. Thus, this analysis gives qualitative information about whether the conserved 3' UTR of *atlas* in *D. melanogaster* is expressed in males in other species. The + or – symbol below each species name indicates whether the top (+) or bottom (-) strand DNA sequence matches the sequence of expressed, 3' UTR mRNA in *D. melanogaster*. Expression of this region was further assessed for some species by RT-PCR; see Fig. S9.

**Table S2. Predicted biochemical properties and levels of conservation for Atlas orthologs identified across *Drosophila* species.**

| Species | Length (a.a.) | Isoelectric Point (pI) | Pairwise BLASTP bit score and e-value with Dmel | Note(s) |
|---|---|---|---|---|
| *D. melanogaster* | 172 | 10.7 | 347, 3e-129 | |
| *D. simulans* | 172 | 10.6 | 281, 1e-103 | |
| *D. sechellia* | 172 | 10.6 | 293, 7e-108 | |
| *D. yakuba* | 178 | 11.4 | 219, 9e-79 | |
| *D. erecta* | 174 | 11.0 | 210, 2e-75 | |
| *D. ananassae* | 170 | 10.3 | 78, 2e-23 | |
| *D. bipectinata* | 233 | 10.3 | 62, 9e-11 | Likely contains an ortholog, given significant BLAST hit and RNAseq evidence of male-specific expression, but gene boundaries are unclear. The 5' end of the ORF contains either ~100 extra codons or an intron, neither of which matches the *atlas* gene structure. Statistics reported at left are for the gene model that includes an intron, GenBank accession XP_017110044.1. |
| *D. elegans* | 171 | 10.4 | 142, 2e-48 | |
| *D. ficusphila* | 180 | 10.6 | 147, 2e-50 | |
| *D. eugracilis* | 176 | 10.9 | 138, 8e-47 | |
| *D. kikkawai* | No ortholog detected at mel or ana syntenic region | | | |
| *D. serrata* | No ortholog detected at mel or ana syntenic region | | | |
| *D. rhopaloa* | 176 | 10.7 | 149, 4e-51 | |
| *D. biarmipes* | 185 | 10.4 | 158, 1e-54 | |
| *D. suzukii* | 179 | 10.6 | 182, 5e-64 | |
| *D. takahashii* | 198 | 10.6 | 165, 4e-57 | |
| *D. obscura* | No ortholog detected | | | |
| *D. miranda* | No ortholog detected | | | |
| *D. persimilis* | >198 | ~10.3 | n.s. | Found in syntenic region to *D. virilis* ortholog, but BLAST results do not support orthology; gap in *D. persimilis* assembly cuts off 3' end of predicted ORF, so length and pI values are estimates |
| *D. pseudoobscura* | 232 | 10.1 | n.s. | |
| *D. willistoni* | No ortholog detected | | | |
| *D. busckii* | 88 | 5.1 | 34.3, 4e-8 | Found in syntenic region to *D. virilis* ortholog, but low pI does not support homologous function |
| *D. hydei* | No ortholog detected | | | |
| *D. arizonae* | No ortholog detected | | | |
| *D. mojavensis* | No ortholog detected | | | |

| | | | |
|---|---|---|---|
| *D. navojoa* | No ortholog detected | | |
| *D. virilis* | 116 | 10.1 | 40, 1e-9 |
| *D. grimshawi* | No ortholog detected | | |

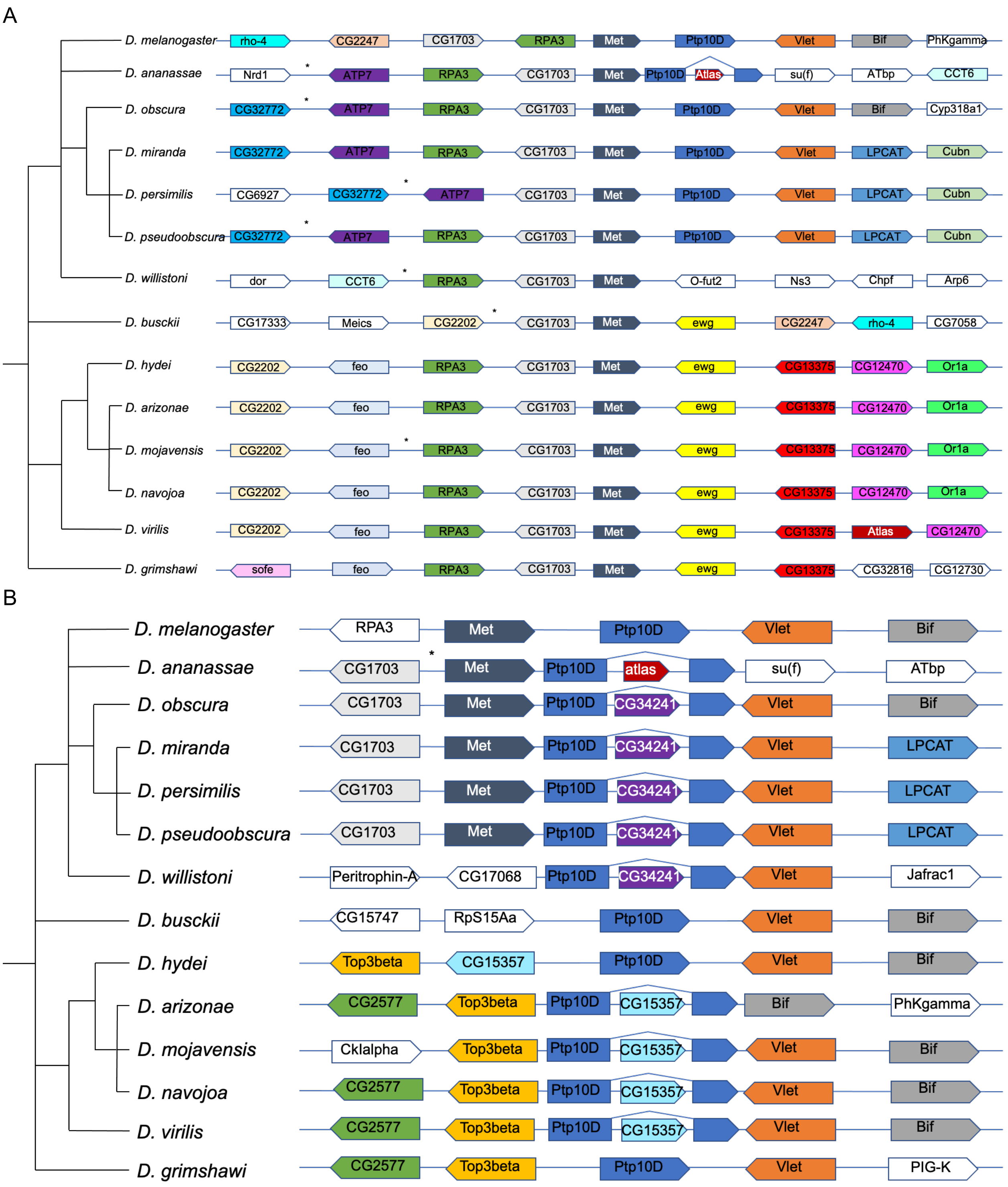

**Fig. S9: Synteny analysis across *Drosophila* species of regions of Muller element A that contain *atlas* in *D. ananassae* and *D. virilis*.** A) *Atlas* is found downstream of the *Met* gene ortholog in both *D. ananassae* and *D. virilis*. This region was therefore searched in multiple additional species. While the genomes of several species harbored unannotated genes in this general region that showed RNA-seq evidence of male expression, all such predicted genes encoded proteins that were significant BLASTP hits to *D. melanogaster* proteins other than Atlas. B) In *D. ananassae*, *atlas* is found in the middle of an intron of the *Ptp10D* gene. However, in some species, *Ptp10D* is no longer syntenic with *Met.* Therefore, we searched for *atlas* orthologs in and around *Ptp10D* across the same set of species. While orthologs of two other *Drosophila* gene have become inserted into a *Ptp10D* intron in other lineages, no additional *atlas* orthologs were found. In both panels, asterisks indicate unannotated genes supported by RNA-seq evidence that were confirmed with BLASTP to be homologs of genes other than *atlas*.

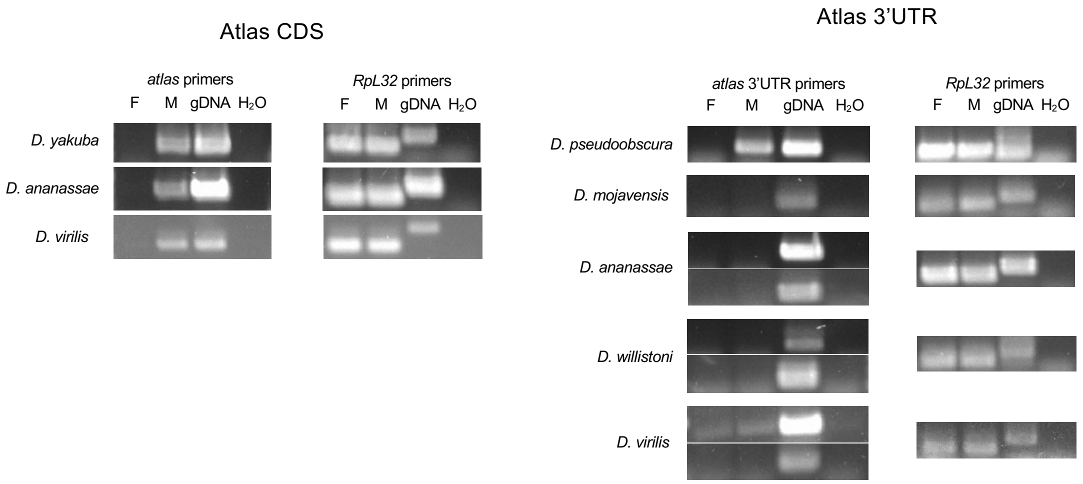

**Fig. S10: RT-PCR of the *atlas* coding sequence (CDS) and non-coding exon (3'UTR) in other *Drosophila* species.** cDNA was prepared from whole males or whole females and analyzed with either *atlas* coding sequence primers, primers designed to a portion of the non-coding exon, or housekeeping gene *RpL32* as a control. The protein-coding region is expressed in a male-specific manner in *D. ananassae* and *D. virilis*, consistent with available RNA-seq data. The non-coding region shows robust male-specific expression in *D. pseudoobscura*, but was not detectable in *D. mojavensis* (one primer pair attempted), *D. ananassae* (two primer pairs attempted) or *D. willistoni* (two primer pairs attempted). One of two primer pairs attempted gave faint, non-sex-specific amplification in *D. virilis*.

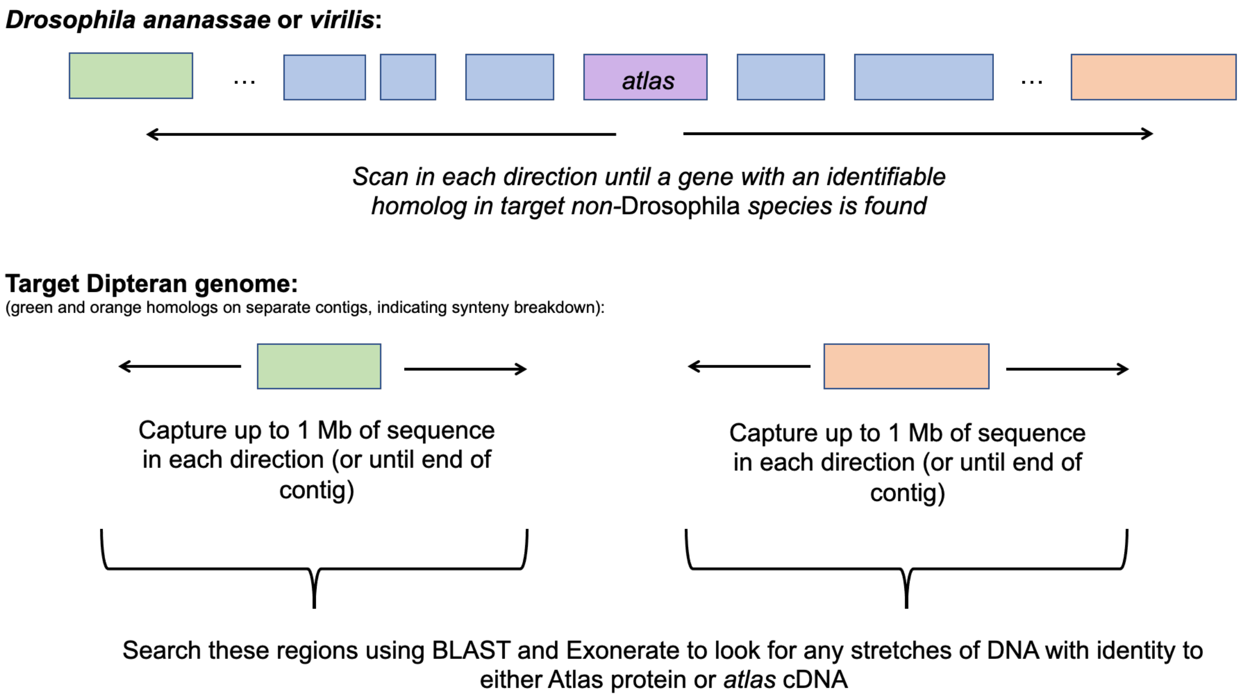

**Fig. S11: Synteny-based method for searching other Dipteran genomes for potential *atlas* orthologs.**

**Table S3. Sources of RNAi lines, degrees of knockdown observed, and hairpins cloned to generate TRiP-style RNAi lines.**

[see included .xlsx file]

**Table S4. Primers used for constructing *atlas*-HA genomic rescue construct and *atlas*-GFP donor template construct.**

***atlas*-HA**
Atlas Rescue F1: GGCATGTCGACCTCGAGTACCCGGGAGCTCGAATTCTAGAcctgaaagcagaacattgta
Atlas Rescue R1: ATGGGTAAAAGATGCGGCCTCCACCGCGGTGGAGATCCATctgaaacgggtccacatcca
Atlas Rescue F2: ATGGATCTCCACCGCGGTGGAGGCCGCA
Atlas Rescue R2: AATTACCACAGTACCTACAATATATTTCCAACACACATCCtcacgtggaccggtgtccgc
Atlas Rescue F3: ggatgtgtgttggaaatatattgtaggtactgtggtaattgtttaacttctcgtttacaaaatg
Atlas Rescue F4: ATTGCCGGCGATATCGGATCCACCGGTGCCTAGGCGCGCCgcgaccgcgcacaaactcat

***atlas*-GFP**
*Cloning of* atlas *CDS into pENTR prior to recombination with pTWG to form atlas-GFP:*
pENTR-atl-F: caccATGGGACGCAAAGGCCACAAG
pENTR-atl-R-nostop: CTGAAACGGGTCCACATCCATG

*Amplification of homology arms for Golden Gate Assembly of scarless CRISPR HDR plasmid:*
Left arm F: cgtctcaggacATGGGACGCAAAGGCCACAAG
Left arm R:
cgtctcaTTAAACAATTACCACAGTACCTACAATATATTTCCAACACACATCgTCACGTGGACCGGTGCTTGTAC
Right arm F: cgtctcaagggTTAACTTCTCGTTTACAAAATGCCC
Right arm R: cgtctcagcatTTGGCGTGGGACTCATTTTGGC
